# Anti-Cancer Immune Priming with Beta-Radioligand Therapy and Isoform-Selective Targeting of 4Ig-B7-H3

**DOI:** 10.1101/2024.12.20.629740

**Authors:** Sarah E. Glazer, Margie N. Sutton, Ping Yang, Federica Pisaneschi, Manu Sebastian, Seth T. Gammon, David Piwnica-Worms

## Abstract

Radioligand therapy (RLT), a re-emerging oncologic strategy using molecularly-targeted therapeutic radioisotopes, clinically reduces tumor burden and enhances survival for select patients with otherwise unresponsive advanced prostate cancer and neuroendocrine tumors. Developing new approaches to next generation targets and a better understanding of systemic immune effects could broaden the impact of RLT. Aside from contributions to immune checkpoint, B7-H3 (*CD276*) is an attractive oncologic target because of its widespread and high differential expression across a variety of solid tumors compared to normal tissues. However, B7-H3 has two isoforms: a 4Ig-B7-H3 isoform, the dominant transmembrane protein expressed on tumors and tumor immune microenvironments (TIME), and a 2Ig-B7-H3 isoform, a soluble ectodomain protein, representing a circulating, and in the context of RLT, significant shed (decoy) antigen. To enhance tumor-specific binding and circumvent confounding soluble 2Ig-B7-H3, a novel IgG2a monoclonal antibody (MIL33B) was generated with high affinity for 4Ig-B7-H3 (72 picomolar) and 8- to 18-fold selectivity over soluble 2Ig-B7-H3. Live cell fluorescence microscopy using AF594-labeled MIL33B demonstrated strong membranous localization and target specificity. PET-CT imaging with ^89^Zr-labeled MIL33B confirmed robust tumor-selective target binding *in vivo* in murine xenograft (HeLa cervical) and syngeneic tumor models (4T1 breast, B16F10 melanoma, and CT26 colorectal) expressing human 4Ig-B7-H3. As a *single* dose beta-emitting systemic RLT therapeutic, ^90^Y-labeled MIL33B (100 μCi) produced 53% long-term survival in a 4Ig-B7-H3-dependent manner in an otherwise fatal established CT26 colorectal tumor model. Immunologic analysis showed that ^90^Y-MIL33B RLT functioned as an immune priming event, engaging downstream CD8^+^ T-cell activation and inducing immunological memory *in vivo*, thus illustrating the potential of systemic beta-RLT to target both primary and metastatic sites. Thus, MIL33B showcases a strategy to selectively target 4Ig-B7-H3 for beta-RLT, warranting further investigation as an immune priming tactic alone or in combination for cancer therapy.

**Statement of significance:** Modest antigen expression levels, even if target tissue-selective, combined with ectodomain shedding (soluble decoy antigens) can generally hinder targeted diagnostic and therapeutic strategies, but are especially challenging for radioligand therapy, PET imaging, and *in vivo* diagnostics wherein high specific activity radioisotopes necessitate use of low masses of biocarrier. Binding, absorption and non-specific tissue deposition of radiolabeled biocarriers by decoy antigens can significantly misdirect systemic radiation, reducing therapeutic efficacy. An antibody development process with a focus aimed at on-target affinity for folded proteins on live cells resulted in a novel picomolar affinity antibody selectively targeting membranous 4Ig-B7-H3 over soluble decoy 2Ig-B7-H3. This antibody shows promise as a transformative systemic beta-radioligand therapy platform for immune priming applications in oncology, and potentially in cardiology, rheumatology, and autoimmunity.

## Introduction

Theranostics is a re-emerging and expanding medical field based on therapeutic interventions directly coupled to imaging to verify the presence and location of a molecular target and assessing disease state ^1^. Radiotheranostics, also referred to as radioligand therapy (RLT), leverages radioisotopes to both image the patient and treat the tumor ^1^. Fundamentally, RLT is systemic therapy whereby the goal is to selectively bind and eliminate primary tumor as well as distant metastases and microscopic disease by delivering molecularly-targeted radioactivity through a single therapeutic intervention **(Figure 1A).** By comparison, conventional external beam radiation therapy (EBRT) targets with curative intent a defined radiation field localized to the primary tumor, and in some applications, to oligometastatic disease ^2^, but many patients present with distant macro- and micro-metastases and continuously shed detectable levels of circulating tumor cells into the systemic circulation outside the radiation field ^2^. Both RLT and EBRT are conventionally thought to achieve their effect through direct radiation-induced toxicity to the tumor cell compartment via irreparable DNA damage, while standard EBRT has been shown to evoke complex immune responses, with evidence for both immune activating (abscopal effect) and immune suppressive actions ^2–4^.

**Figure 1.**
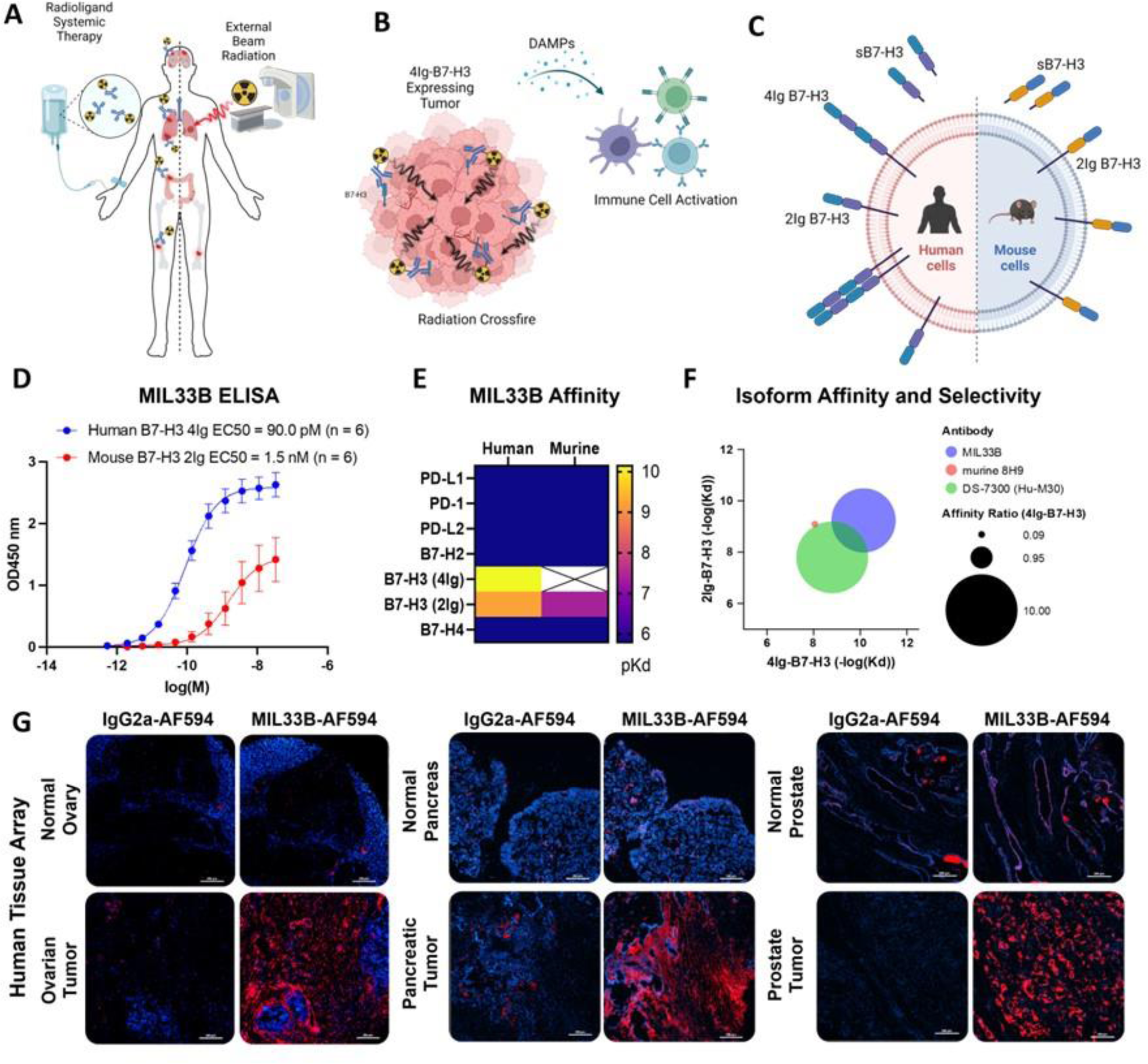
Development of a High Affinity, 4Ig-Isoform Selective Antibody for Beta-Radioligand Therapy. **A)** Radioligand therapy (RLT) represents systemic therapy whereby the goal is to selectively bind and eliminate primary tumor sites as well as distant metastases and microscopic disease by delivering molecularly-targeted radioactivity through a single therapeutic intervention as compared to conventional external beam radiation therapy (EBRT) which targets with curative intent a defined radiation field localized to the primary tumor, and in some applications, to oligometastatic disease. **B)** Schematic describing the role “Crossfire” whereby utilizing a 4Ig-B7-H3 molecularly targeted beta-radioligand therapy, the longer distance of the beta-particle deposition and the heterogenous expression of 4Ig-B7-H3 on the tumor and tumor microenvironement allow for a larger, yet still targeted deposition of radioisotope. Upon damage and tumor cell killing, damage-associated molecular patterns (DAMPs) may be released and provide immune priming signals. **C)** B7-H3 (CD276) is a type-I transmembrane immune modulatory protein with two isoforms found in humans (4Ig-B7-H3 and 2Ig-B7-H3) and a single isoform found in mice (2Ig-B7-H3). There is a soluble 2Ig-component that can be found in circulation, which may serve as a “sink” for non-differentiated isoform-specific anti-B7-H3 antibodies. **D)** ELISA of serial dilutions of MIL33B from different production lots demonstrated high affinity to plate-bound extracellular domain of human 4Ig-B7-H3 (EC50 = 90.0 pM, 95% CI: 62 pM – 130 pM; n = 6), and lower affinity to plate-bound extracellular domain of mouse 2Ig-B7-H3 (EC50 = 1.7nM, 95% CI: 0.47 nM – 5.0 nM; n = 6). **E)** Heatmap of biolayer interferometry analysis of MIL33B binding to B7-H3 homologues and common immune checkpoint proteins. MIL33B had the highest affinity for human 4Ig-B7-H3 and selection over the 2Ig-B7-H3 human isoform. **F)** MIL33B demonstrated greater isoform affinity and selectivity compared to other anti-B7-H3 antibodies. **G)** Immunofluorescence staining of fresh frozen human tissue microarrays with normal and paired tumor tissue. Images represent IgG2a-AF594 labeled isotype control (left, red) and MIL33B-AF594 (right, red) were overlaid on DAPI (nuclear DNA, blue) for normal (top) and cancerous tissue (bottom). Scale bars represent 200 µm.

The concept of RLT has been developing for more than 70 years. Exemplary early instances include using different forms of radioactive iodine to diagnose (e.g., ¹²⁴I) and treat (e.g., ¹³¹I) thyroid cancers ^1^. Later, clinical success with ¹³¹I-tositumomab (Bexxar), a CD20-directed radiotherapeutic antibody indicated for the treatment of patients with CD20-positive refractory non-Hodgkin’s lymphoma, and recent US Food and Drug Administration (FDA) approvals of ¹⁷⁷Lu-dotatate (Lutathera; Novartis) targeting SSTR2-positive neuroendocrine tumors, and ¹⁷⁷Lu-PSMA-617 (Pluvicto; Novartis) for PSMA-positive prostate cancer, have raised RLT into the mainstream of cancer care options for selected patient populations. Overall, RLT is thought to produce local radiation-induced reactive oxygen species (ROS), single/double strand DNA breaks and oxidative stress, thereby damaging targeted cancer cells and the nearby TIME ^1, 5–7^. However, the field is hampered by a lack of detailed understanding of mechanisms of action of this new class of anti-cancer therapeutics and the narrow scope of FDA-approved targets, which portend an unmet medical need to expand the repertoire of second-generation targets and high-affinity ligands.

Modern RLT agents are modular, built from a targeting ligand, a linker, and radioisotope. Radioisotopes can be used to deliver highly potent alpha-particles, beta-particles or auger-electrons, each with unique properties and ranges of tissue damage that can be leveraged depending on the targeting context. Beta-particles are negatively charged, and deposit less energy (0.2 KeV/mm linear energy transfer (LET)) compared to alpha-particles or auger-electrons, but over a longer distance (0.05-12 mm) ^8^. In the context of molecular-targeted RLT, this longer distance energy deposition of beta-particles allows for “crossfire” and cell killing even in the face of genetic heterogeneity of the tumor and/or tumor immune microenvironment **(Figure 1B)**. Not every cell needs to express the molecular target for beta-RLT to be effective ^8^. Furthermore, careful selection of isotope or isotope pairs enables either pre-imaging of therapy or post-therapy imaging to readily localize tumors and metastases, quantify pharmacokinetics of the ligand in various tissues, and/or provide multi-scalar biomarkers for either patient selection or patient management.

In this regard, B7-H3 (CD276), a homologue of the B7 superfamily of cell-surface immune co-regulatory proteins, is found in a wide variety of cancers (colon, renal, cervical, esophageal, hepatocellular, neuroblastoma, breast, pancreatic, prostate, head and neck, glioblastoma, ovarian, and NSCLC, among others) ^9–17^, correlating with poor prognosis, reduced survival, and increased metastasis ^18^. While *B7-H3* mRNA is readily detected in most human tissues, by contrast, B7-H3 protein expression is essentially non-existent in normal tissues, being tightly regulated by microRNAs, particularly miR-29 ^19^. B7-H3 protein is present in only a subset of normal cells and tissues, such as placenta and testis, wherein the extracellular domain of B7-H3 would be poorly accessible to a targeting agent in the blood ^20, 21^. Note that B7-H3 protein can be induced on multiple components of the TIME (cancer cells, cancer-associated vasculature, neutrophils, monocytes, dendritic cells and other antigen-presenting cells ^22–28^), and thus, combined with low protein content in normal tissue, B7-H3 has attracted interest as an oncologic target in recent years ^9, 29^ and may provide an ideal target for RLT.

Importantly, human B7-H3 is a type 1 transmembrane protein expressed in two isoforms, a 4Ig-B7-H3 isoform consisting of an extracellular ectodomain comprising two pairs of immunoglobulin constant and variable domains (IgC/IgV, IgC/IgV) ^22, 30–32^, and a 2Ig-B7-H3 isoform containing only one extracellular IgC/IgV pair ^33^. This stands in contrast to rodents, wherein only the 2Ig isoform is expressed ^32^, and may be responsible for the species-specific differential roles of B7-H3 in regulating immune checkpoint observed in early studies ^34^. In humans, a soluble component (s2Ig-B7-H3), likely representing ectodomain shedding of B7-H3, similar to other type 1 and 2 membrane proteins and members of the B7 family ^35, 36^, has been identified and accumulates in compartments outside the TIME, such as the blood pool and cerebral spinal fluid [39, ^37–39^. Increased circulating s2Ig-B7-H3 has been identified in the serum of patients during progression of malignancies and inflammatory states ^40–44^. This is of particular concern for the development of imaging and low-mass RLT therapeutics because s2Ig-B7-H3 can act as a decoy target and ligand sink, potentially diminishing efficacy and increasing toxicities of ligands that target both isoforms. None-the-less, abundant evidence indicates that 4Ig-B7-H3 is the dominant isoform present on the cell surface of human cancers ^22, 30–32, 45^, and unique to B7 family members, the presence of 4Ig and 2Ig isoforms may provide a “handle” to develop a 4Ig-tumor-selective ligand, such as an antibody for RLT to overcome challenges presented by circulating B7 isoforms.

Herein, a next-generation targeted beta-radioligand antibody therapeutic was extensively validated *in vitro* and *in vivo* yielding functional cures in xenograft and syngeneic immunocompetent murine tumor models. The biochemical targeting mechanism was firmly established, and the high cure rate in a syngeneic model opened the opportunity to study systemic immune responses. Mechanistically, 4Ig-B7-H3-targeted beta-RLT functioned as a primary immune priming event shown to engage downstream CD8^+^ T-cell activation and induce immunological memory *in vivo*, thus experimentally demonstrating the potential systemic engagement of the immune system during targeted beta-RLT.

## Results

### Development of a High Affinity, 4Ig-Isoform Selective Antibody for Beta-Radioligand Therapy

To develop a 4Ig-B7-H3-specific antibody with both high affinity to the folded extracellular domain of the human 4Ig isoform and selectivity over the s2Ig isoform, New Zealand White (NZBWF1/J) or Balb/C mice were immunized by foot pad injection with murine L-cells transduced with cassettes expressing human 4Ig-B7-H3 on the cell surface. Host sera were screened by ELISA to assess binding to the extracellular domains of both human 4Ig-B7-H3 and murine 2Ig-B7-H3, and candidate hybridomas subsequently scaled for further evaluation. Candidates were counter-screened by live cell fluorescence binding assays to HeLa cells, known to express 4Ig-B7-H3 and secrete s2Ig-B7-H3 ^46^, yielding 3 preliminary antibody candidates that could bind to folded, glycosylated human 4Ig-B7-H3 in the face of s2Ig-B7-H3 *in cellulo*. Final prioritization of leads was ultimately based on high affinity binding by ELISA to folded human 4Ig-B7-H3 **(Figure 1E)**. Subsequently, hybridoma expansion was performed to further evaluate murine monoclonal antibody candidate 33B, hereafter known as Molecular Imaging Laboratory 33B (MIL33B), isotype IgG2a, for affinity and specificity against various B7 family members, animal species and isoforms.

Affinity to the folded ectodomain of human 4Ig-B7-H3 in an ELISA format yielded an EC_50_ of 90.0 pM (95% CI: 62 pM -130 pM; n =6), substantially higher affinity (18-fold) compared to the plate-bound extracellular domain of murine 2Ig-B7-H3 (EC_50_ = 1.7 nM; 95% CI: 0.47 nM – 5.0 nM; n = 6) **(Figure 1D, Supplemental Figure 1)**. Next, MIL33B binding kinetics were measured for folded human, murine and porcine B7-H3 isoforms by biolayer interferometry as a model of binding selectivity between the 4Ig extracellular domain and s2Ig in circulation, resulting in calculated K_D_ values of 7.23×10^−11^ M for human 4Ig-B7-H3, 5.80×10^−10^ M for human 2Ig-B7-H3, 4.11×10^−8^ M for mouse 2Ig-B7-H3, and 1.02×10^−10^ M for porcine 4Ig-B7-H3 **(Table 1**, **Figure 1E, Supplemental Figure 2)**. These profiles for MIL33B demonstrated significantly higher affinity for 4Ig-B7-H3 when compared to other B7-H3 antibodies that have been tested clinically, e.g., 8H9 and MGA271 ^47–51^, with 8-fold selectivity for the human 4Ig isoform compared to the human 2Ig isoform **(Table 1**, **Figure 1F)**, thus, presenting the opportunity to enhance tumor-targeting by traversing soluble decoy in circulation.

**Table 1.**
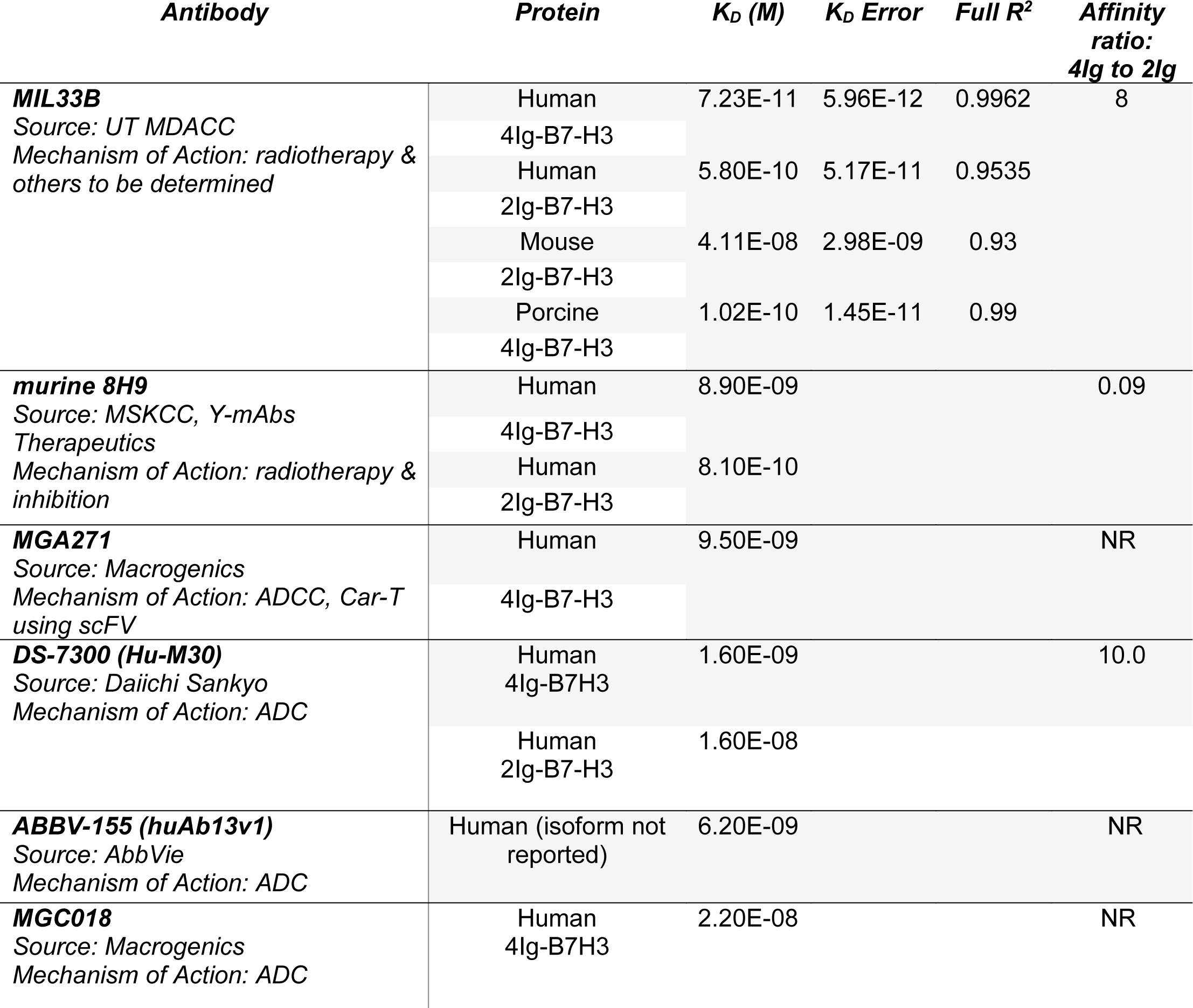
Comparative Affinity and Selectivity of MIL33B to B7-H3 Isoforms. Biolayer interferometry analysis of MIL33B binding to the extracellular domains of human 4Ig-B7-H3, human 2Ig-B7-H3, mouse 2Ig-B7-H3, and porcine 4Ig-B7-H3. MIL33B demonstrated higher affinity and higher selectivity for the human 4Ig isoform over the 2Ig isoform when compared to other anti-B7-H3 antibodies that have been tested clinically (NR = not reported or calculable). Affinities from other antibodies collected from published values ^29, 47, 48, 132, 133^. ADC = antibody-drug conjugate, ADCC =antibody dependent cellular cytotoxicity, UT MDACC= University of Texas MD Anderson Cancer Center, MSKCC = Memorial Sloan Kettering Cancer Center.

The specificity of MIL33B for B7-H3 versus B7 family members was further evaluated by determination of the affinity of MIL33B to other human and mouse B7 family members and common immune checkpoint proteins. K_D_ values by ELISA of MIL33B against plate-bound mouse and human B7-H2, B7-H4, PD-L1, PD-L2, PD-1, and CTLA-4 were not measurable (>1×10^−6^ M) **(Figure 1E)**, yielding > 5-orders of magnitude selectivity. Similarly, biolayer interferometry analysis of the binding of MIL33B to human and mouse B7-H2, B7-H4, B7-H1(PD-L1), PD-L2, and PD-1 were also below the limit of detection demonstrating orders of magnitude of selectivity **(Table 2 and Supplementary** Figure 3).

**Table 2.** MIL33B Affinity to Human and Mouse B7 Immunoglobulin Superfamily Members. Biolayer interferometry analysis of MIL33B binding to B7-H3 homologues and common immune checkpoint proteins demonstrated no significant affinity, confirming MIL33B specificity for multi-species B7-H3.

| *Protein* | *KD(M)* | *KD Error* | *Full R^2* |
| --- | --- | --- | --- |
| **Human PD-L1** | > 1.6E-6 | N/A | 0 |
| **Mouse PD-L1** | > 1.6E-6 | 1.15E-07 | 0 |
| **Human PD-1** | > 1.6E-6 | N/A | 0 |
| **Mouse PD-1** | > 1.6E-6 | N/A | 0 |
| **Human PD-L2** | > 1.6E-6 | 1.24E-08 | 0 |
| **Mouse PD-L2** | > 1.6E-6 | 3.77E-08 | 0 |
| **Human B7-H2** | > 1.6E-6 | 4.52E-09 | 0 |
| **Mouse B7-H2** | > 1.6E-6 | 1.0E-12 | 0 |
| **Human B7-H4** | > 1.6E-6 | 4.19E-09 | 0 |
| **Mouse B7-H4** | > 1.6E-6 | 2.21E-11 | 0 |

### Validation of Selective Binding of CDRs of MIL33B to Human Cancers versus Normal Tissues

To evaluate tumor-differentiated binding of B7-H3 by MIL33B, fixed-frozen human tissue arrays containing microarrayed tissue from multiple organ sites of “normal” or “tumor” tissue were stained using MIL33B labeled with AF594 or the appropriate IgG2a-AF594 labeled control antibody. Differential and tumor-specific binding was observed using MIL33B-AF594 for several tumor types including ovarian, pancreatic, and prostate tumors **(Figure 1G)**.

### Validation of CDR Binding and 4Ig-B7-H3 Protein Targeting by Live Cell Fluorescence Microscopy

To evaluate B7-H3 targeting of MIL33B *in cellulo* (and later *in vivo*), several high and low 4Ig-B7-H3-expressing human and mouse cancer cell lines were generated. By Western blot analysis, parental HeLa cells (HeLa^+/+^) showed relatively high endogenous expression of human 4Ig-B7-H3 and low 2Ig-B7-H3 **(Supplemental Figure 4A)**, while a clonal HeLa B7-H3 CRISPR/Cas9 knockout cell line (HeLa^−/−^; KO) showed no detectable protein ^52^. As expected, 4T1 (murine triple negative breast cancer), B16F10 (murine melanoma cell lines), and CT26 (murine colon cancer) demonstrated low minimal expression of the murine 2Ig-B7-H3 isoform relative to human tumor cells **(Supplemental Figure 4A**). Note that the commercial antibody leveraged in the Western blot experiments likely overestimated the levels of rodent versus human protein as its primary cognate antigen was mouse 2Ig with cross-reactivity to the human protein. To develop murine lines containing the human 4Ig-isoform of B7-H3, 4T1, B16F10, and CT26 cells were batch transfected with human 4Ig-B7-H3 lentiviral particles, generating cells expressing human 4Ig-B7-H3 protein at levels comparable or less than to those observed in human cancer cells **(Supplemental Figure 4A)**.

Next, binding *in cellulo* of MIL33B and specificity for human 4Ig-B7-H3 were tested. MIL33B and a matched non-targeted isotype control antibody (IgG2a) were labeled with Alexa594. HeLa 4Ig-B7-H3^+/+^, HeLa B7-H3^−/−^, 4T1 4Ig-B7-H3, 4T1 negative vector, B16F10 4Ig-B7-H3, B16F10 negative vector, CT26 4Ig-B7-H3, and CT26 negative vector cells were incubated with either MIL33B-Alexa549, mouse IgG2a-Alexa549, or no antibody. Cell-associated antibody was assessed by live cell fluorescence microscopy, which enabled evaluation of both membrane association and internalization of antibodies over time. MIL33B demonstrated high cell-specific binding to endogenously high 4Ig-B7-H3-expressing human cells (HeLa^+/+^) and cells transduced with human 4Ig-B7-H3 (CT26 4Ig-B7-H3, 4T1 4Ig-B7-H3 and B16F10 4Ig-B7-H3) with a prominent plasma membrane pattern and evidence of internalization by 40 min to 1 hr. MIL33B selectivity for 4Ig-B7-H3 versus other tumor surface antigens, FcR binding, and macropinocytosis was demonstrated via the vastly superior binding to cells expressing 4Ig-B7-H3 versus various negative controls, including B7-H3 Hela KO cells, negative vector control murine cells (reflecting low endogenous murine 2Ig-B7-H3), cells incubated with mouse IgG2a-Alexa549, and autofluorescence of unlabeled cells alone **(Figure 2A-F, Supplemental Figure 4B-C)**. Of note, we observed heterogeneous 4Ig-B7-H3-specific cell labeling of MIL33B in murine cell lines transduced with human 4Ig-B7-H3, consistent with expectations for cells not clonally selected for high 4Ig-B7-H3 expression after transfection. Further specificity of MIL33B versus non-specific membrane adhesion and macropinocytosis was independently validated by pre-incubating HeLa cells with excess un-labeled MIL33B, which blocked MIL33B-Alexa549 binding as accessed by live cell fluorescence microscopy **(Figure 2B)**. In addition, because the MIL33B isotype was mouse IgG2a, which has high affinity to mouse FcγR ^53^, additional validation of specificity versus FcγR binding was evaluated. Indeed, MIL33B pre-blocking was far superior to IgG2a pre-blocking, indicting no substantial Fc-mediated binding to solid tumor-expressed Fc receptor family, consistent with the previously demonstrated superiority of MIL33B-AF594 compared to murine IgG2a-AF594 live cell binding and retention **(Figure 1C)**. Overall, MIL33B showed high specificity for human 4Ig-B7-H3 protein versus other tumor antigens, tumor expression of FcγR, non-selective membrane binding, and macropinocytosis *in cellulo*.

**Figure 2.**
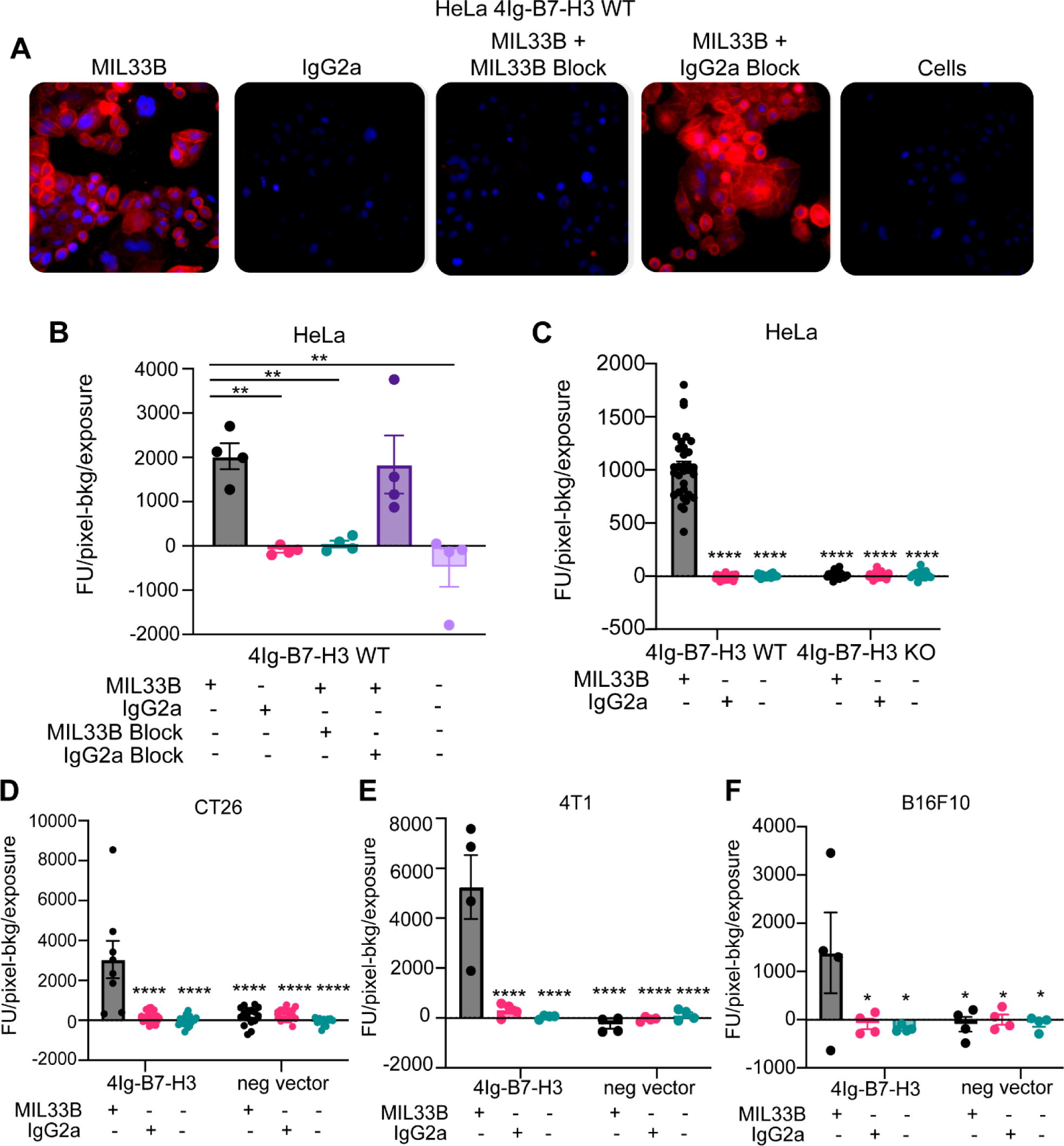
MIL33B has high affinity and specificity for the human 4Ig-B7-H3 isoform. A. Immunofluorescence live-cell staining of multiple 4Ig-B7-H3 expressing cell lines demonstrated 4Ig-B7-H3 specific binding and membrane localization of MIL33B. High endogenous 4Ig-B7-H3-expressing human HeLa cells incubated with Alexa549-labeled MIL33B demonstrated high intensity membrane-specific localization compared to Alexa549-labeled IgG2a or cells alone when incubated for 1 hr. Pre-incubation with un-labeled MIL33B was superior to murine IgG2a in displacing Alexa549-labeled MIL33B (cold block). **B-F.** Quantification of MIL33B-AF594 or IgG2a-AF594 staining intensity (Fluorescence Units-Background/Seconds) following 1 hour incubation with antibodies at 37°C and 5% CO_2_ humidity. HeLa WT **(B)**, HeLa WT and KO **(C)**, CT26 **(D)**, 4T1 **(E)**, or B16F10 **(F)** were cultured under normal conditions.

### Evaluation of MIL33B Targeting Tumor Cell 4Ig-B7-H3 Protein In Vivo and Application to Immuno-PET

Next, to determine if MIL33B could be utilized as an effective 4Ig-B7-H3 targeting antibody *in vivo* and demonstrate utility for radio-immuno PET applications, ^89^Zr-DFO conjugates of MIL33B and isotype control (^89^Zr-DFO-MIL33B and ^89^Zr-DFO-IgG2a, respectively) were generated and characterized. Live animal PET imaging allowed characterization of tumor-specific net retention, kinetics, and whole body biodistribution of MIL33B in different tumor models, and relative organ-specific clearance over time. Zirconium-89 was selected to exploit the long isotope half-life (3.3 days), which both aligned with the long half-life of circulating antibodies *in vivo* and provided ample time to conduct longitudinal imaging of mice injected with each radiolabeled antibody. Similar antibody conjugates have also been translated into humans for various PET applications ^54^.

To synthesize ^89^Zr-DFO-MIL33B and ^89^Zr-DFO-IgG2a, each antibody was first conjugated to the chelator DFO **(Supplemental Figure 5A)**. LC/MS analysis of DFO-MIL33B and DFO-IgG2a identified a mass shift consistent with the addition of two DFO chelators on average per antibody **(Supplemental Figure 5D, 5G)**. Subsequently, DFO-MIL33B and DFO-IgG2a were labeled with ^89^Zr-oxalate, purified on a PD-10 column, and characterized by radio-TLC and radio-SEC-HPLC. ^89^Zr-DFO-MIL33B labeled with 62.7% ± 13.7% (n = 4) chelation efficiency as measured by radio-TLC. Radio-SEC-HPLC of ^89^Zr-DFO-MIL33B immediately after purification demonstrated 82% purity and specific activity of 16.6 μCi/μg (6.14×10^−4^ GBq/μg) (n = 1) **(Supplemental Figure 5B-C)**. ^89^Zr-DFO-IgG2a labeled with 54.5% ± 14.2% (n = 4) chelation efficiency as measured by radio-TLC. Radio-SEC-HPLC of ^89^Zr-DFO-IgG2a immediately after purification demonstrated 87% purity and specific activity of 10 μCi/μg (3.7×10^−4^ GBq/μg) (n = 1) **(Supplemental Figure 5E-F).**

After intravenous injection of either ^89^Zr-DFO-MIL33B or ^89^Zr-DFO-IgG2a, the biodistribution and 4Ig-B7-H3-specific tumor-targeting in nude mice harboring either HeLa 4Ig-B7-H3^+/+^ tumors or HeLa B7-H3^−/−^ tumors were evaluated by PET-CT at 24, 72, and 144 hours post injection. Furthermore, to evaluate the displaceability of ^89^Zr-DFO-MIL33B *in vivo,* some mice received 200 μg i.v. of non-radiolabeled MIL33B (cold block) 1 hr before receiving ^89^Zr-DFO-MIL33B.

HeLa cells were again selected as a key test case because they endogenously express 4Ig-B7-H3, secrete s2IgB7-H3, and are infiltrated in the murine immune environment *in vivo* with MDSCs expressing FcγR ^55, 56^. ^89^Zr-DFO-MIL33B binding in the tumor compartment in mice harboring HeLa 4Ig-B7-H3^+/+^ tumors (4.27 ± 1.28 %ID/cc at 24 hours, 5.37 ± 1.40 %ID/cc at 72 hours, and 5.02 ± 1.30 %ID/cc at 144 hours post tracer injection; n = 3) was greater at all timepoints compared to mice harboring HeLa B7-H3^−/−^ tumors (1.77 ± 0.63 %ID/cc at 24 hours, 1.69 ± 0.68 %ID/cc at 72 hours, and 1.67 ± 0.61 %ID/cc at 144 hours post tracer injection; n = 3) **(Supplemental Figure 6A, E, I, L)**. Note that both ^89^Zr-DFO-isotype control and pre-blocking experiments exhibited significantly different blood pool clearance times necessitating individualized normalization to blood pool to test for 4Ig-B7-H3 specific tumor retention. Herein, blood normalized tumor retention in HeLa 4Ig-B7-H3^+/+^ tumors imaged with ^89^Zr-DFO-MIL33B (3.25 ± 0.65; n = 3) was significantly higher compared to HeLa B7-H3^−/−^ tumors (0.79 ± 0.13; n = 3; * p = 0.0201, un-paired two-tailed t-test), HeLa B7-H3^+/+^ tumors imaged with ^89^Zr-DFO-MIL33B after pre-blocking with cold MIL33B (1.02 ± 0.18; n = 3; * p = 0.0292, un-paired two-tailed t-test), HeLa 4Ig-B7-H3^+/+^ tumors imaged with ^89^Zr-DFO-IgG2a (0.77 ± 0.15; n = 3; * p = 0.0201, un-paired two-tailed t-test), or HeLa B7-H3^−/−^ tumors imaged with ^89^Zr-DFO-IgG2a (0.67 ± 0.13; n = 3; * p = 0.0173, un-paired two-tailed t-test) **(Figure 3A-B)**. These results demonstrated that ^89^Zr-DFO-MIL33B was specific for human 4Ig-B7-H3 in the tumor compartment *in vivo* versus normal tissues or other tumor antigens, and demonstrated the requirement for 4Ig-B7-H3 antigen-specific CDRs for tumor retention.

**Figure 3:**
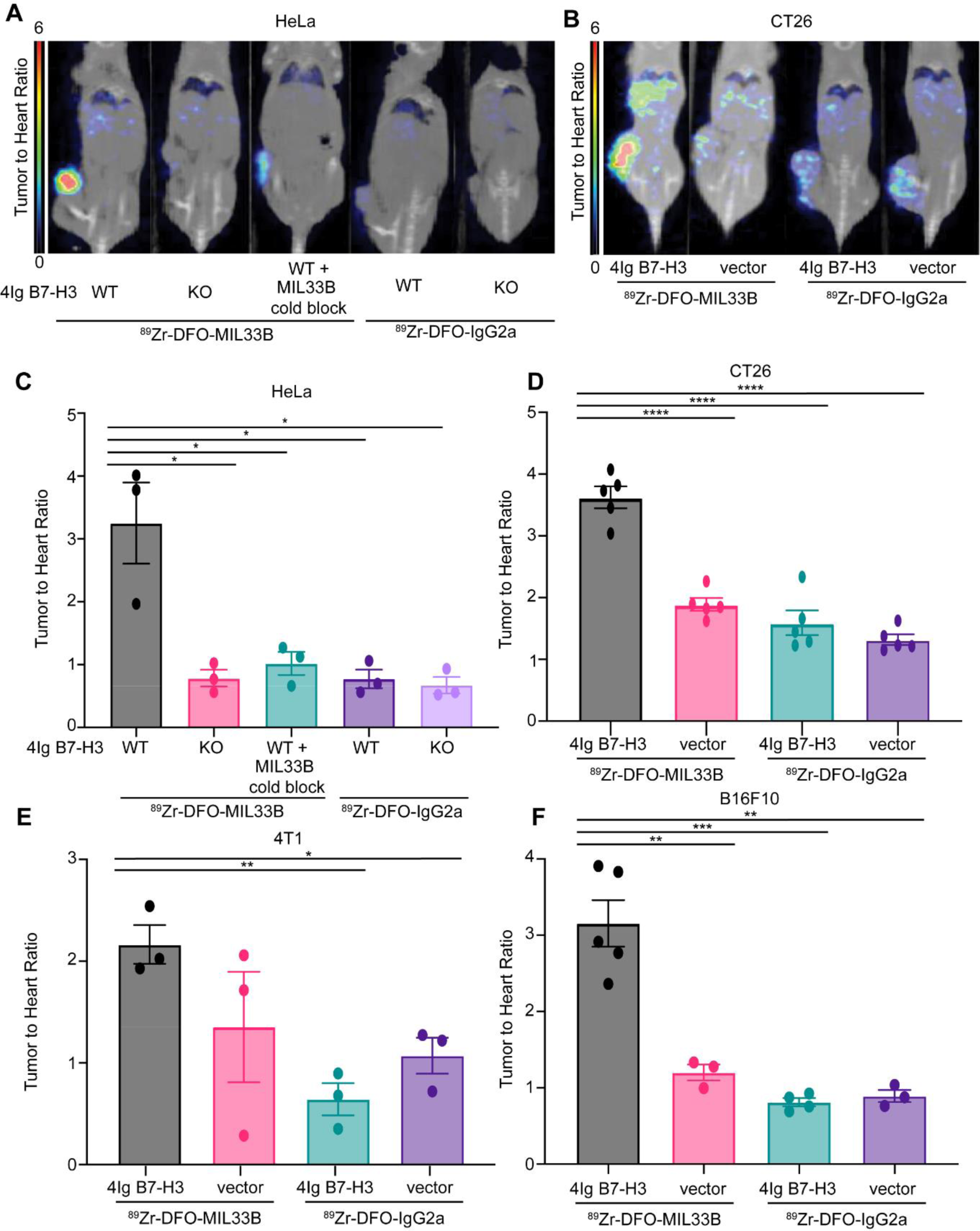
Immuno-PET demonstrates MIL33B targeting to tumor-specific 4Ig-B7-H3 in vivo. PET/CT images, SUV normalized to heart to correct for animal-to-animal variability in clearance between genotypes for HeLa xenografts (A) and CT26 syngenic tumor models (B). Images acquired at 72 hours post-injection revealed significantly higher binding and retention of 89Zr-DFO-MIL33B in mice harboring HeLa B7-H3 +/+ tumors compared to HeLa B7-H3−/− tumors or HeLa B7-H3+/+ tumors pre-treated with cold MIL33B, or HeLa B7-H3+/+ tumors and HeLa B7-H3−/− tumors imaged with 89Zr-DFO-IgG2a (A). Similar on-target (B7-H3 expressing tumor) uptake was observed using the syngeneic mouse model for colorectal cancer, CT26 (B). Normalized SUV of 89Zr-DFO-MIL33B was significantly retained within 4Ig-B7-H3 expressing tumors compared to genotype negative controls or using isotype control, 89Zr-DFO-IgG2a. The phenotype was observed in four different models including the human cervical cancer cell line, HeLa (C), as well as several murine syngeneic models transduced with human 4Ig-B7-H3 compared to respective tumors transfected with negative vector control or imaged with 89Zr-DFO-IgG2a; CT26 (D), triple negative breast cancer model 4T1 (E), melanoma model B16F10 (F). Comparisons are determined by un-paired two tailed t-tests, * = p < 0.05, ** = p < 0.01, *** p < 0.001.

Robustness was further validated by testing 4Ig-B7-H3-expressing 4T1, CT26, and B16F10 syngeneic tumor models. These tumors represent two highly immune infiltrated models and one “immune desert” model (B16F10), respectively, and furthermore enabled assessment of targeting across TH1 (C57BL/6) and TH2 (Balb/c) murine backgrounds ^57^. Balb/c mice were implanted subcutaneously with 4T1 4Ig-B7-H3 cells on the right flank and 4T1 negative vector cells on the left flank, and subsequently injected i.v. with either ^89^Zr-DFO-MIL33B or ^89^Zr-DFO-IgG2a prior to imaging by PET-CT at 24, 72, and 144 hours post tracer injection. Similar to the HeLa tumor model, statistically significant greater retention of ^89^Zr-DFO-MIL33B was observed in 4T1 4Ig-B7-H3 tumors compared to 4T1 negative vector tumors, or retention of ^89^Zr-DFO-IgG2a in 4T1 4Ig-B7-H3 tumors, or ^89^Zr-DFO-IgG2a in 4T1 in neg vector tumors **(Supplemental Figure 6E-H).** When tumor-specific retention at 72 hours post tracer injection was normalized to blood pool (heart-associated) counts and compared across tumor types and antibodies, significantly higher normalized retention of ^89^Zr-DFO-MIL33B in 4T1 4Ig-B7-H3 tumors (2.16 ± 0.19) was observed compared to the retention of ^89^Zr-DFO-IgG2a in 4T1 4Ig-B7-H3 tumors (0.64 ± 0.16, ** p 0.0036, un-paired two-tailed t-test) or in 4T1 neg vector tumors (1.07 ± 0.18, * p = 0.0135, un-paired t-test) **(Figure 3C-D)**. Although a decrease in the average normalized tumor retention of ^89^Zr-DFO-MIL33B in 4T1 negative vector tumors was seen compared to 4T1 4Ig-B7-H3 tumors, the effect size was small. Nonetheless, these results demonstrated overall that expression of human 4Ig-B7-H3 in 4T1 tumors significantly increased tumor-specific retention of ^89^Zr-DFO-MIL33B *in vivo* relative to isotype control, and CDR-specific binding to the tumor compartment in a highly immune infiltrated environment, replete with FcγR expression and internalizing MDSCs^56, 58^.

Next, the biodistribution and tumor specific-retention of ^89^Zr-DFO-MIL33B and ^89^Zr-DFO-IgG2a in mice harboring B16F10 (immune desert) tumors either transduced with human 4Ig-B7-H3 or transfected with the relevant negative vector control was evaluated. Statistically significant greater tumor retention of ^89^Zr-DFO-MIL33B in B16F10 4Ig-B7-H3 tumors was again observed compared to B16F10 negative vector tumors, or tumor retention of ^89^Zr-DFO-IgG2a in B16F10 4Ig-B7-H3 tumors or B16F10 negative vector tumors **(Supplemental Figure 6I-K)**. When blood pool normalized, there was significantly higher normalized tumor retention of ^89^Zr-DFO-MIL33B in B16F10 4Ig-B7-H3 tumors (3.16 ± 0.30; n = 5) compared to B16F10 negative vector tumors (1.20 ± 0.69; n = 3; ** p = 0.0032, un-paired two-tailed t-test), retention of ^89^Zr-DFO-IgG2a in B16F10 4Ig-B7-H3 tumors (0.81 ± 0.41; n = 4; *** p = 0.0003, un-paired two-tailed t-test) or in B16F10 neg vector tumors (0.89 ± 0.52; n = 3; ** p = 0.0015, un-paired two-tailed t-test). These results demonstrated that expression of human 4Ig-B7-H3 in the B16F10 murine melanoma model significantly increased tumor-specific retention of MIL33B *in vivo* **(Figure 3E-F)**.

Finally, the biodistribution and tumor-specific retention of ^89^Zr-DFO-MIL33B and ^89^Zr-DFO-IgG2a in the CT26 murine colon cancer model was investigated. Mice were implanted subcutaneously on the right flank with either CT26 4Ig-B7-H3 cells or CT26 negative vector cells and imaged by PET-CT at 24 and 72 hours post tracer injection (i.v.). As before, following blood pool normalization, significantly higher tumor-specific retention of ^89^Zr-DFO-MIL33B was found in CT26 4Ig-B7-H3 tumors (3.62 ± 1.62; n = 5) compared to CT26 negative vector tumors (1.89 ± 0.85; n = 5; *** p < 0.0001, un-paired two-tailed t-test), or tumor retention of ^89^Zr-DFO-IgG2a in CT26 4Ig-B7-H3 tumors (1.59 ± 0.71; n = 5; *** p < 0.0001, un-paired two-tailed t-test) or in CT26 negative vector tumors (1.32 ± 0.59, tumors; n = 5; *** p < 0.0001, un-paired two-tailed t-test) **(Figure 3G-H, Supplemental Figure 6)**. Again, in this model, these results demonstrated that expression of human 4Ig-B7-H3 significantly increased tumor-specific retention of MIL33B *in vivo*.

### MIL33B as a 4Ig-B7-H3-Targeted Radio-Ligand Therapeutic

After validating *in vivo* the 4Ig-B7-H3-specific tumor-targeting properties of MIL33B by PET-CT, MIL33B was engineered into an antibody radio-conjugate for the treatment of solid tumors. The overarching treatment strategy was to chelate MIL33B to yttrium-90, a radionuclide used clinically in other radio-immunotherapies that possesses a long-range beta emission (11.3 mm) allowing cell “crossfire” within heterogeneous TIMEs ^59–61^ **(Figure 1B)**.

MIL33B was first conjugated to a bifunctional chelator containing DOTA, followed by chelation with yttrium-90 **(Figure 4A, Supplemental Figure 7)**. LC/MS of DOTA-MIL33B was consistent with a mixed pool of one to zero DOTA chelators per antibody **(Supplemental Figure 7C-D)**. MIL33B-DOTA labeled with yttrium-90 at greater than 61.9% ± 17.2% (n = 4) chelation efficiency as determined by radio-TLC. Radio-SEC-HPLC of purified ^90^Y-DOTA-MIL33B demonstrated > 99% purity (n = 5) and a specific activity of 3.15 ± 0.50 μCi/μg (1.17×10^−4^ GBq/μg) (n = 4) **(Figure 4B and Supplemental Figure 7A-B)**.

**Figure 4.**
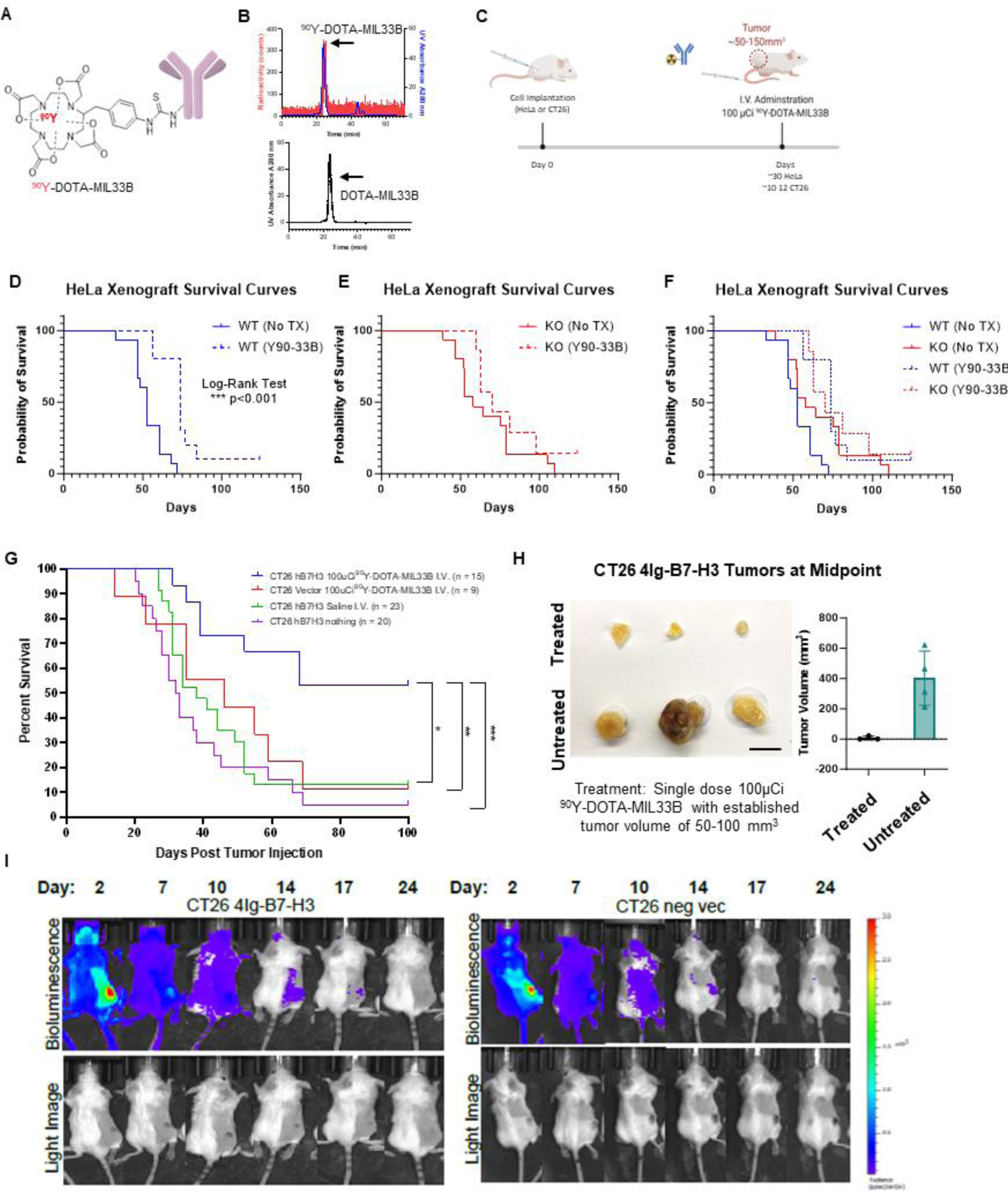
Therapeutic Efficacy of I.V. ^90^Y-DOTA-MIL33B *in Vivo*. **A.** Diagram of ^90^Y-DOTA-MIL33B. **B.** Radio-TLC was used to document the labeling efficiency of ^90^Y-DOTA-MIL33B. **C.** Experimental schema used for *in vivo* studies to deliver a single dose of 100 μCi ^90^Y-DOTA-MIL33B or saline for tumor treatment. **D.** 5 x 10^6^ HeLa WT cells were injected into the right flank of athymic nude mice and allowed to establish a tumor for 30 days prior to treatment of half of the mice with ^90^Y-DOTA-MIL33B. Survival curves for untreated (solid blue) or treated (dashed blue) were plotted. Single dose treatment with ^90^Y-DOTA-MIL33B significantly extended survival *** p< 0.001. **E.** 5 x 10^6^ HeLa KO cells were injected into the right flank of athymic nude mice and allowed to establish a tumor for 30 days prior to treatment of half of the mice with ^90^Y-DOTA-MIL33B. Survival curves for untreated (solid red) or treated (dashed red) were plotted. **F.** Kaplan-Meier survival curves for WT and KO HeLa xenografts were plotted demonstrating the right shift in survival following knockout of B7-H3 from the tumor compartment, and further right shifting with anti-B7-H3 targeted radioligand therapy, ^90^Y-DOTA-MIL33B. **G.** CT26 (vector control or hB7-H3 transduced) tumors were implanted subcutaneously in XX mice and allowed to establish to a size between 50-100 mm^3^ (10-12 days) prior to treatment with saline or a single dose of ^90^Y-DOTA-MIL33B (100 μCi). Kaplan-Meier survival curves were plotted for CT26-hB7-H3 tumors untreated (purple), CT26-hB7-H3 tumors treated with i.v. saline (green), CT26-vector tumors treated with ^90^Y-DOTA-MIL33B (100 μCi) (red) CT26-hB7-H3 tumors treated with ^90^Y-DOTA-MIL33B (100 μCi) (blue). 4Ig-B7-H3 expressing CT26 tumors treated with a single dose of ^90^Y-DOTA-MIL33B (100 μCi) had significantly prolonged survival compared to all other treatment groups. **H.** Midpoint tumors collected 7 days post treatment with 100 μCi ^90^Y-DOTA-MIL33B for three mice in each group were excised and photographed. Tumor volume for all midpoint tumors was plotted for CT26-hB7-H3 that were treated vs. untreated (N=4 in each group). **I.** Cherenkov imaging of ^90^Y-DOTA-MIL33B. Mice harboring CT26 hB7-H3 tumors and CT26-vector tumors were imaged on an IVIS spectrum 2, 7, 10, 14, 17 and 24 after I.V. administration of 100 µCi ^90^Y-DOTA-MIL33B. Measurement of Cherenkov radiation demonstrated tumor uptake in both CT26-hB7-H3 and CT26-vector tumors that diminished overtime.

### Efficacy of Beta-RLT on 4Ig-B7-H3-Expressing HeLa Tumor Xenografts In Vivo

As with fluorescence and ^89^Zr-PET imaging, RLT efficacy was first tested with HeLa tumors to generate preliminary signals of efficacy in a model that secretes soluble B7-H3 **(Supplemental Figure 8)**. Mice harboring HeLa 4Ig-B7-H3^+/+^ tumors (n = 10) or mice harboring HeLa B7-H3^−/−^ KO tumors (n = 7) received a single dose (100 µCi) of ^90^Y-DOTA-MIL33B 30 days after tumor implantation and followed for 120 days **(Figure 4C, Supplemental Figure 9)**. Compared with untreated animals harboring HeLa WT and KO xenografts, single dose ^90^Y-DOTA-MIL33B right-shifted median survival for all animals treated with 4Ig-B7-H3-RLT **(Figure 4D-F)**, where a genotype specific further right shift was observed for WT mice, where known expression of 4Ig-B7-H3 was present on the tumor xenograft (log-rank test (*** P<0.001)); Hazard Ratio ^90^Y-DOTA-MIL33B/No Tx: 0.2706, 95%CI ratio: 0.1140 to 0.6423) **(Figure 4D)**. The smaller, yet present right shift in the survival for the KO bearing tumors, although not statistically significant, may be explained by the beneficial effect observed in other murine and human studies where radiation to the liver, the know excretion organ for antibodies and ^90^Y-DOTA-MIL33B, has been demonstrated in animal models and human studies to provide anti-tumor effects ^62^ **(Figure 4E)**. When all survival curves are superimposed **(Figure 4F)**, the magnitude of response by a single dose RLT can be appreciated wherein ^90^Y-DOTA-MIL33B-RLT extended survival, even producing cures in this immunocompromised model, and beyond that observed from knockout of the target (B7-H3) alone. Modified interrupted time series analysis was conducted to compare individual growth rates 15 days before treatment and 15 days after treatment on a per tumor basis. There was a significant decrease in tumor growth rates in treated HeLa 4Ig-B7-H3^+/+^ tumors (2-way ANOVA, ** p = 0.0019) versus HeLa B7-H3^−/−^ KO tumors (2-way ANOVA, p = 0.1027), demonstrating an initial treatment response to ^90^Y-DOTA-MIL33B in this immunocompromised model **(Supplemental Figure 9C-D).** Knockout of B7-H3 in HeLa cells substantially shifts their growth rates compared to wild type cells, the mechanism by which has been dissected by Sutton et. al. ^63, 64^. However, long term, there were no differences in overall survival at the end point of treatment **(Supplemental Figure 9A-B)**, likely due to intrinsic differences in tumor growth rates that depend upon B7-H3 and an eventual rebound in tumor growth as observed in this immunocompromised model.

### Efficacy of Beta-RLT in a Syngeneic Tumor Model In Vivo

Next, syngeneic immune competent models of RLT were tested. To fully evaluate the therapeutic efficacy of intravenous ^90^Y-DOTA-MIL33B, mouse cohorts harboring established CT26 4Ig-B7-H3 or negative vector tumors were tumor-size selected (50-150 mm^3^) 10 to 12 days after cell implantation **(Figure 4C)**. Untreated mice were followed regardless of initial tumor size. Based on the results of a pilot dose-response trial, a single dose (100 µCi) of ^90^Y-DOTA-MIL33B i.v. was characterized in detail for 4Ig-B7-H3-based therapeutic responses. Thus, over a combination of five independent experimental cohorts, 15 mice harboring CT26 4Ig-B7-H3 tumors received ^90^Y-DOTA-MIL33B i.v., 9 mice harboring CT26 negative vector tumors received ^90^Y-DOTA-MIL33B i.v., 23 mice harboring CT26 4Ig-B7-H3 tumors received an equivalent volume of saline i.v., and 20 mice harboring CT26 4Ig-B7-H3 tumors received no treatment. In the CT26 4Ig-B7-H3 group that received a *single* dose of 100 µCi ^90^Y-DOTA-MIL33B i.v., an initial tumor regression of 11 out of 15 tumors was observed; however, 3 tumors emerged at later time points. In the CT26 negative vector group that received 100 µCi ^90^Y-DOTA-MIL33B i.v., an initial tumor regression of 2 out of 9 tumors was observed and 1 of these tumors later emerged. In the CT26 4Ig-B7-H3 group that received intravenous saline, tumor regression of only 3 out of 23 tumors was found, and in the CT26 4Ig-B7-H3 groups that did not receive any treatment, spontaneous tumor regression of 1 out of 20 tumors was observed **(Supplemental Figure 10A-D)**. When mice were followed for 100 days, 53% long-term survivors were observed in mice harboring CT26 4Ig-B7-H3 tumors treated with 100 µCi ^90^Y-DOTA-MIL33B i.v., compared to only 11% long-term survivors in mice harboring CT26 negative vector tumors receiving the same dose (* p = 0.0376, log-rank test), 13% long-term survivors in mice harboring CT26 4Ig-B7-H3 tumors treated with saline (** p = 0.0034, log-rank test), and 5% long-term survivors in mice harboring CT26 4Ig-B7-H3 tumors that received no treatment (*** p = 0.0003, log-rank test) **(Figure 4G)**. When tumors in a subset of animals were harvested at the typical *mid-point* of treatment (day 6-8 post initial ^90^Y-DOTA-MIL33B injection), representative tumors from mice harboring CT26 4Ig-B7-H3 tumors treated with ^90^Y-DOTA-MIL33B were significantly smaller compared to untreated mice harboring the same tumor cells **(Figure 4H).**

The low yield positron emission of ^90^Y ^65^ is below the threshold for detection by conventional animal PET imaging at the dilute, yet therapeutic, concentrations of ^90^Y tested herein ^66^. As an alternative to PET imaging, Cherenkov radiation emitted by the beta decay was observed by optical imaging to non-invasively qualify net retention of ^90^Y-DOTA-MIL33B in the tumor compartment longitudinally, which confirmed preferential localization of ^90^Y-DOTA-MIL33B to 4Ig-B7-H3 tumors versus control tumors *in vivo* **(Figure 4I)**, similar to that observed by PET-CT with ^89^Zr-labeled MIL33B. Furthermore, white light mode imaging allowed observation of individual tumors over time, confirming complete regression of representative CT26 4Ig-B7-H3 tumors and continued growth of CT26 negative vector tumors **(Figure 4I).** In addition to complete tumor regression, no obvious radiotoxicity was observed in mice that responded to treatment with ^90^Y-DOTA-MIL33B (weight, fur, diet consumption, and activity were grossly normal as monitored both by the investigators and impartial veterinary staff). These results demonstrated that a single i.v. injection of ^90^Y-DOTA-MIL33B led to complete regression of over half of established CT26 murine colorectal tumors in a 4Ig-B7-H3-dependent manner, demonstrating promise for a high therapeutic index.

### ^90^Y-DOTA-MIL33B Engages Adaptive Immunity

After establishing that 100 µCi of ^90^Y-DOTA-MIL33B i.v. led to differential regression of established CT26 tumors that expressed human 4Ig-B7-H3 compared to CT26 tumors with low B7-H3 expression, the mechanism of action of this radio-therapeutic response was assessed. First, the composition of the TIME in mice treated with ^90^Y-DOTA-MIL33B was characterized by histology. Tumors from mice harboring CT26 4Ig-B7-H3 tumors that had received 100 µCi of ^90^Y-DOTA-MIL33B i.v. were excised at the midpoint of treatment, fixed and saved for histology after radioactive decay **(Figure 5A)**. Representative IHC tumor staining for CD3^+^, CD4^+^, and CD8^+^ antigens demonstrated positive staining for each of these markers inside the tumor compartment, indicating that CD8^+^ and CD4^+^ T cells had widely infiltrated into the TIME in mice treated with ^90^Y-DOTA-MIL33B i.v. **(Figure 5A).** In addition, further histologic quantification indicated that there were CD8^+^ T-cells present in the tumor compartment, and as expected with a model of heterogenous target expression, there were heterogenous distributions of T-cells on a per animal basis. Similarly, the CD3^+^, CD8^+^, and CD4^+^ cell populations tended to co-correlate across groups (**Supplemental Figure 11**).

**Figure 5.**
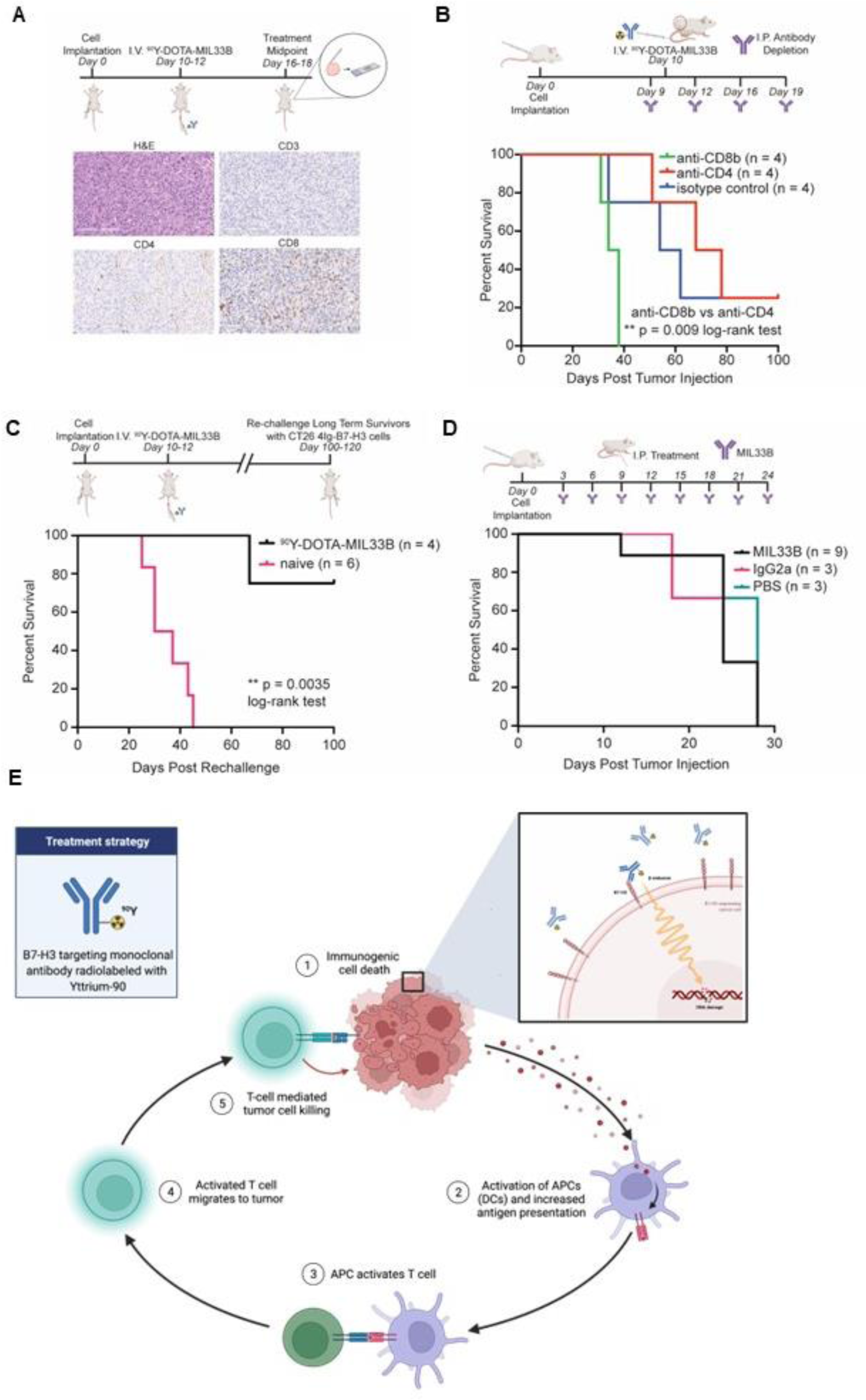
Treatment with ^90^Y-DOTA-MIL33B Elicits and Immunological Response. **A.** Representative histology of a ^90^Y-DOTA-MIL33B treated tumor. Mice harboring established CT26 hB7-H3 tumors were treated with 100 μCi i.v. ^90^Y-DOTA-MIL33B. Six days post treatment mice were sacrificed and tumors were fixed in formalin and subsequently stored in 70% ethanol and allowed to decay. Histological staining demonstrated positive CD3, CD4 and CD8 staining in the tumor compartment indicating that both CD4 and CD8 T cells are present. Scale bars represent 200 μm. **B.** Immune cell depletion studies identify CD8b cells as essential for ^90^Y-DOTA-MIL33B response. Mice harboring CT26 hB7-H3 tumors were treated with 100 µCi of ^90^Y-DOTA-MIL33B i.v. One day before treatment and 2, 4, and 7 days after treatment mice received 250ug I.P. of either anti-CD4 or anti-CD8b depleting antibodies or the relevant isotype control antibody, rat isotype control. Survival curves were compared for isotype control (blue), anti-CD4 (red) and anti-CD8b (blue) using the log-rank test. (** = p < 0.01). **C.** Re-challenge of ^90^Y-DOTA-MIL33B i.v. responsive mice. Mice harboring CT26-hB7-H3 tumors that were treated with 100 µCi of ^90^Y-DOTA-MIL33B i.v. and whose tumors regressed were rechallenged with CT26-hB7-H3 cells on the opposite flank 100-120 days after the initial tumor implantation. 75% of rechallenge mice did not re-grow tumors compared to naïve mice who all grew tumors and reached endpoint in response to cell implantation. Survival curves were compared by the log-rank test (** = p < 0.01). **D.** MIL33B monotherapy alone does not support survival advantage as a monotherapy. Mice implanted with 1e5 CT26-hB7-H3 cells received either 200ug I.P. of MIL33B (black), mouse IgG2a isotype control antibody (pink) or PBS (teal) every three days till endpoint. Kaplan-Meier survival curves were plotted for each treatment group. Single agent treatment with MIL33B did not produce a survival benefit for CT26-hB7-H3 tumor-bearing mice. **E.** Working model of the immunological response elicited by treatment of CT26-hB7-H3 tumors with ^90^Y-DOTA-MIL33B, beta-radioligand therapy. Beta emissions induce cell damage, leading to immunogenic cell death, release of DAMPs and engagement of innate immunity (1). DAMPs activate APCs (2) leading to increased antigen presentation and activation of T cells (3). T cells then migrate to the tumor (4) and exert additional cytotoxic effects on tumor cells (5).

Having validated antibody binding to tumor-associated 4Ig-B7-H3, mechanistic intervention studies were conducted. Because CD8^+^ and CD4^+^ T cells were present in the TIME in mice that had previously received 100 µCi of ^90^Y-DOTA-MIL33B i.v., the dependence of treatment response on components of the immune system downstream of the initial radio-ablation was further evaluated. Herein, mice harboring CT26 4Ig-B7-H3 tumors received intravenous treatment with 100 µCi of ^90^Y-DOTA-MIL33B as described previously. However, one day before treatment, mice received 250 µg i.p. of either anti-CD4- or anti-CD8b-depleting antibodies or the relevant isotype control antibody (anti-rat isotype antibody). Mice subsequently received i.p. injections of depleting antibodies or relevant control antibodies on day 2, day 6, and day 9 post treatment **(Figure 5B)**. For mice that received anti-CD4-depleting antibodies, 3 out of 4 tumors initially regressed, but later, 2 of those 3 tumors reemerged. In mice that received anti-CD8b-depleting antibodies, no tumors regressed, and in mice that received anti-rat isotype control antibodies 1 out of 4 tumors regressed **(Supplemental Figure 12A-C)**. When mice were followed for 100 days post tumor implantation, no mice in the group that received anti-CD8b-depleting antibody plus 100 µCi of ^90^Y-DOTA-MIL33B were deemed long-term survivors and these mice reached endpoint faster than mice that received anti-CD4-depleting antibodies plus 100 µCi of ^90^Y-DOTA-MIL33B (25% long-term survivors; ** p = 0.0091, log-rank test). In addition, injections of rat isotype control did not affect the therapeutic efficacy of 100 µCi of ^90^Y-DOTA-MIL33B (25% long-term survivors) **(Figure 5B).** These results indicated that CD8b+-expressing cells, but not CD4+-expressing cells, were necessary to confer long-term therapeutic effects of ^90^Y-DOTA-MIL33B *in vivo* and pointed to CD8+ T cells or CD8+ NKT cells as essential contributors to long-term survival.

### Documenting Immunological Memory

While both the HeLa and CT26 models required human 4Ig-B7-H3 in the tumor compartment for beta-RLT to induce an early decrease in tumor volumes, long term cures were primarily evident in the immune competent CT26 system. In the immunocompromised nude mouse model there was a single long term survivor in the WT 90Y-DOTA-MIL33B group vs 0 in the untreated group. While not statistically detectable this is consistent with a rank ordering of the long-term survivors with immunocompromised nu/nu mice vs syngeneic mice. These data and the observed RLT-induced tumor immune infiltrate suggested a potent mechanistic role for cellular adaptive immunity. Thus, mice harboring CT26 4Ig-B7-H3 tumors previously treated with 100 µCi of ^90^Y-DOTA-MIL33B and deemed long-term survivors (90 to 110 days post treatment), received a second re-challenge of fresh CT26 4Ig-B7-H3 cells implanted on the left (opposite) flank. Re-challenged mice were compared to naïve mice (never before inoculated with tumor cells nor treated) also implanted subcutaneously with matched fresh CT26 4Ig-B7-H3 cells on the left flank **(Figure 5C)**. Both cohorts were followed for 100 days post tumor cell implantation. At study end point, in a combination of two independent experiments, all naïve mice had grown tumors and reached terminal endpoint, while only 1 out of the 4 re-challenged mice grew a tumor (** p = 0.0035, log-rank test); the remaining 3 re-challenged mice were able to fend off tumor re-challenge and survive **(Figure 5C)**. These results indicated that mice with CT26 4Ig-B7-H3 tumors that had previously responded to radio-ligand therapy developed immunological memory, a signature indicating that adaptive immunity was indeed engaged for long-term survival following treatment with ^90^Y-DOTA-MIL33B.

### Non-Radiolabeled (Cold) MIL33B Therapy

As an additional control, MIL33B efficacy as a single agent antibody was explored at a mass excess to radiotherapy in a manner consistent with established immunotherapy regimens for other B7-family targeted anti-cancer therapies. This experiment served as a control to estimate the potential therapeutic contribution of binding of cold MIL33B to human 4Ig-B7-H3, acting either through blockade of B7-H3 signaling pathways, complement, ADCP, or ADCC. Mice implanted with CT26 4Ig-B7-H3 cells received either 200 μg of MIL33B (non-radiolabeled) i.p., 200 μg of mouse IgG2a isotype control i.p., or PBS i.p., every three days post cell implantation until endpoint **(Figure 5D)**, a typical regimen for immune checkpoint therapy in pre-clinical models ^67^. Note that immune checkpoint regimens involve substantially higher therapeutic mass of MIL33B (> 32-fold) than the mass used for a single administration of ^90^Y-DOTA-MIL33B (≤ 50 μg). Treatment with cold MIL33B was far inferior to ^90^Y-DOTA-MIL33B therapy, i.e., showing no observable effect in this tumor model **(Figure 5D and Supplemental Figure 13A-C).** These results further supported the proposed model wherein the effects of ^90^Y-DOTA-MIL33B were due to 4Ig-B7-H3-mediated retention of a therapeutic radionuclide, yttrium-90, within the local tumor environment that initiated immunogenic cell death, leading to engagement of innate immunity, an immune priming cascade, and ultimately activation of adaptive immunity **(Figure 5E).**

## Discussion

While B7-H3 expression is generally thought to be “pro-tumor” and immunosuppressive in humans, the detailed signaling functions of the 4Ig and 2Ig isoforms of B7-H3 remain under investigation with both co-stimulatory and co-suppressive immune effects being reported both in cancer and in other inflammatory disorders ^33, 68–82^. Indeed, the complexities may be species-specific as rodents have evolutionarily lost the 4Ig isoform. Thus far, non-isoform selective anti-B7-H3 antibody binding or blocking strategies have met with limited clinical success in oncology, particularly relative to anti-B7 therapies targeting the PD-1/PD-L1 axis, although combination anti-B7-H3 therapy with anti-PD-L1 in the context of *PTEN*/*TP53-*deficiency shows promise in pre-clinical prostate cancer and AML models ^83, 84^. Interestingly, soluble ectodomain antigens (in this case, shed ectodomain 2Ig-B7-H3) present challenges to antibody therapy because antibodies bound to antigens are often cleared faster from circulation, altering pharmacokinetics ^46, 85, 86^. Furthermore, antibody antigen complexes can lead to off-target toxicities both in the local micro-environment, and in FcγR2-expressing cells ^87, 88^. These off-tumor effects can be further compounded in the case of B7-H3 because the concentration of circulating s2Ig-B7-H3 can increase with tumor progression or advanced inflammation, dynamically impacting dosing regimens or potential toxicities ^42, 89–93^.

Learning from challenges faced by first generation anti-B7-H3 antibodies, a new high affinity, isoform-selective anti-4Ig-B7-H3 antibody was herein generated through early selective pressure for live cell binding and imaging *in vivo* using strategic models known to present other possible confounders of binding, including possible tumor specific glycosylation ^94, 95^ and dimerization ^63, 64^. Through early use of live cell selection, B_max_ (the maximum specific binding) in addition to K_d_ could also be subtly maximized through the selection for antibody binding epitopes that are accessible in the context of folded proteins, quaternary structure, and cell surface supramolecular architectures. Such stringency would not be available for antibodies focused on blocking interactions that must compete for accessibility to the target against high local concentration of competitors, which by definition lower B_max_, and therefore binding potential in the low-mass regime. Isoform-selective anti-4Ig-B7-H3 binding was characterized and validated through biochemical assays *in vitro*, live cell binding and microscopy assays with knockout, knock-in, and blocking studies *in cellulo*, and similarly executed tumor binding and PET-CT imaging studies *in vivo*, which were robust to multiple cell lines, immune-competent and immune-defective murine models, and using both human cells as well as 4Ig-B7-H3 transfected mouse cells. Non-specific membrane adhesion, macro-pinocytosis, B7 paralogues, tumor surface antigens, tumor-expressed FcγR binding, blood accessible murine surface antigens *in vivo*, FcγR binding *in vivo*, and enhanced permeability-retention (EPR) *in vivo* were tested and demonstrated non-contributory. Finally, through this development process, the platform utility of the antibody for systemic delivery of cargo to 4Ig-B7-H3 compartments *in vivo*, such as the TIME, was demonstrated with PET-CT imaging, Cherenkov imaging, and critically, beta-RLT (radio-theranostics) in a syngeneic CT26 tumor model. Of note, CT26 tumors are characterized as an aggressive MSI-low colorectal carcinoma model radioresistant to conventional external beam radiotherapy [73-75], thus, setting a high bar for pre-clinical advancement of beta-RLT ^96–98^.

Further analysis of the TIME of CT26 4Ig-B7-H3 tumors at the midpoint of beta-RLT with ^90^Y-DOTA-MIL33B revealed that CD4^+^ and CD8^+^ T cells had infiltrated into the TIME. In addition, mice harboring established CT26 4Ig-B7-H3 whose tumors had previously regressed in response to ^90^Y-DOTA-MIL33B developed immunological memory. These phenotypes generally depend upon activation of adaptive immunity. To test this proposed mechanism of immune memory and cure, *in vivo* depletion assays demonstrated that the final therapeutic efficacy of ^90^Y-DOTA-MIL33B depended upon CD8b^+^-expressing cells. These results led to a working model wherein the therapeutic effects of ^90^Y-DOTA-MIL33B rely on early radiation-induced initiation of immunogenic cell death leading to secondary adaptive immune responses and activation of cytotoxic T cells. This is further supported by the early response to treatment observed in an immunocompromised tumor model of mice bearing established human HeLa B7-H3^+/+^ and HeLa B7-H3^−/−^ tumors, wherein treatment with a single dose of ^90^Y-DOTA-MIL33B resulted in a significant initial regression of HeLa B7-H3^+/+^-treated tumors superior to HeLa B7-H3^−/−^ tumors. In addition, survival of ^90^Y-DOTA-MIL33B-treated HeLa B7-H3^+/+^ tumors compared to standard survival of HeLa xenografts showed a significant right-shift in overall survival and an observed long-tail consistent with a model wherein RLT with ^90^Y-DOTA-MIL33B provided an immune priming event. We propose that the often ignored low levels of adaptive immunity present in nude mice enabled some level of therapeutic efficacy and a small subset of long-term survivors in ^90^Y-treated cohorts.

These results point to the general potential of beta-emission-based 4Ig-B7-H3-targeted RLT as immune priming agents. Previous work has implicated two pathways in radiation-induced activation of the adaption immune system, the release of DAMPs and activation of the cGAS-STING pathway ^99, 100^. Both pathways can stimulate antigen presenting cells to present neo-antigens to T cells. Further work with relevant knockout mouse models may pinpoint signaling pathways activated by ^90^Y-DOTA-MIL33B and additional pharmacodynamic biomarkers downstream of target engagement. In this regard, it is interesting to speculate that the cross-fire properties of beta-emitting therapeutic radionuclides such as ^90^Y (and ^177^Lu, ^67^Cu) may enhance the repertoire of neoantigens released within the TIME, since both 4Ig-B7-H3-expressing and nearby non-expressing cells will be impacted by local beta emissions, perhaps overcoming intercellular genetic heterogeneity and resistance mechanisms. In addition, although T cells were present in the TIME after treatment with ^90^Y-DOTA-MIL33B, it was unknown whether ^90^Y-DOTA-MIL33B actively recruited peripheral T cells to the TIME or if ^90^Y-DOTA-MIL33B changed the activation setpoint of T cells already residing in the TIME. Discerning this difference may help to further elucidate the mechanism of action of ^90^Y-DOTA-MIL33B as well as understand the milieu of the TIME most responsive to RLT. Furthermore, it would be predicted that early disease or micro-metastases will also be impacted by beta-RLT, enabling systemic treatment of micro-disease, not just local control of a primary tumor **(Figure 1A-B)**.

Clinically, radioembolization with ^90^Y in the treatment of hepatic malignancies has previously been shown to increase the CD8^+^ T-cell and NKT-cell infiltration into the TIME ^101^. Pre-clinically, others have demonstrated that ^90^Y-based radionuclide therapy can activate the adaptive immune system in a Non-Hodgkin’s Lymphoma model and synergize with immunotherapy to increase CD8^+^ T-cells in the TIME of pre-clinical solid tumor models ^102, 103^. ^90^Y-radio-embolization can elicit an adaptive immune response in humans ^101^, and human anti-mouse antibody reactions can predict survival after radio-immunotherapy in humans ^104–106^. Immune activation downstream of ^90^Y-DOTA-MIL33B treatment point to the clinical potential of 4Ig-B7-H3-targeted RLT as both a stand-alone agent and potential immune priming agent for use in combination with other immune checkpoint or immunotherapies for the treatment of many 4Ig-B7-H3-expressing solid and liquid tumors.

Is the potential immune priming restricted to ^90^Y, or is there evidence of broader differential activation of immunity by beta-radiation, or more broadly, charged particle therapy? Careful interrogation of the clinical survival curves for ^177^Lu-PSMA show not only extensions in median survival, but a “long tail” now recognized as a characteristic of immune activation and engagement of adaptive immunity ^107^. Furthermore, because of its long-term success and the robustness of the agent, lessons derived from the history of thyroid cancer treatment concluding that a simple radioactive salt for radio-iodine therapy (^131^I^−^) as standard of care might be overlooked. External beam radiation was first tried and found to be inferior to ^131^I^−^. Importantly, there are many accessible radioisotopes of iodine that have different emission characteristics, including gamma-emitters, auger-electron emitters, and beta-emitters (^131^I^−^). Through years of clinical optimization for a non-nuclear target, the beta-emitter won out and yields cure rates of >>> 85%, wherein the long survival tail is obscured by its very height. Furthermore, thyroid cancer, like CT26, is an “immune swamp” ^108^. Finally and intriguingly, proton therapy, a charged particle therapy externally delivered also can engender cures of CT26 tumors in contrast to standard external beam X-ray therapy in pre-clinical models ^109^. A recent phase 3 trial also points to potent curative effects of proton therapy, particularly in head and neck *cancer* ^110^.

B7-H3 is under active exploration as an immune checkpoint, ADC, and CAR T-cell target, and has been previously explored as a radio-immunotherapy target. Pre-clinically, ^212^Pb-376.96 has been explored for the treatment of human ovarian cancer xenografts and ^131^I-4H7 has been investigated in the treatment of human renal cell carcinoma xenografts ^111–113^. These pre-clinical studies demonstrated either extended overall survival or tumor growth rate inhibition ^111–113^. However, these pre-clinical studies mainly utilized immunodeficient models, and not immunocompetent models, thus curtailing the full range of mechanism of action or resistance to a radio-immunotherapeutic when both the innate and adaptive immune systems are present. 8H9, a murine anti-B7-H3 antibody, is the only anti-B7-H3 RLT that has been investigated clinically ^47, 114^, focused on non-humanized ^131^I-8H9 for the treatment of metastatic neuroblastoma or medulloblastoma by intrathecal injection. ^131^I-8H9 and ^124^I-8H9 are also under investigation for the treatment of CNS-relapsed rhabdomyosarcoma and DIPG, respectively ^115–119^. These trials have yielded promising results in closed CNS compartments ^115–119^, but attempts at systemic therapy of solid tumors have not been reported, perhaps related to the overall lower affinity of the primary antibody as well as the lack of differential isoform selectivity of 8H9, which has higher affinity for the human 2Ig isoform compared to the 4Ig isoform. The selective 2Ig-B7-H3 affinity may be sufficient when administered in a closed compartment such as the CNS; however, systemic therapy will be substantially hindered by the soluble 2Ig decoy. This is especially important in the context of systemic RLT, wherein high specific activity radiolabeling methods require use of low masses of antibody (compared to masses of antibody used with immune checkpoint or ADCs). Low mass regime therapeutics, such as RLT, utilizing antibodies with poor selectivity will bind blood decoys, form micro- and macro-immune complexes, remain in circulation or deposit in tissues ^120^, and thus, deplete the primary antibody, adversely impact pharmacokinetics, and abrogate therapeutic effectiveness. Indeed, soluble decoys may confound targets relevant to the RLT field, including most B7 family members, PSMA, EGFR, and IGF1R ^121–125^. The promise of pre-clearing and pre-targeting strategies used in RLT to manage decoy targets can be a two-edged sword because of immune complex formation^126^. Traversing soluble decoys through engineered antibodies as demonstrated herein provides an alternative and/or complementary strategy.

In summary, ^90^Y-DOTA-MIL33B is an effective radio-ligand therapeutic for the treatment of 4Ig-B7-H3-expressing solid tumors and may have broad potential both as a stand-alone RLT agent and in combination with other immune modulators. Because MIL33B is a high affinity 4Ig-B7-H3-specific targeting agent engineered to traverse soluble (shed ectodomain) s2Ig-B7-H3 *in vivo*, MIL33B provides a modular platform of CDRs and antibody fragments for the delivery of additional therapeutic cargoes, including other beta-emitting and alpha-emitting isotopes, antibody-drug conjugates, peptides, bi- or tri-specific antibodies, and cell-based therapies in oncology. Additional cell-based therapeutics and applications may apply to cardiovascular disease, inflammation, and rheumatological disorders engaging 4Ig-B7-H3, including immune-depletion of memory cells in autoimmunity ^127, 128^.

## Brief Methods

See **Supplement Materials** for detailed methods.

### Antibody Development and MIL33B Production

Anti-B7-H3 antibodies were generated at the MDACC Monoclonal Antibody Core Facility (protocol 00000620-RN01 through RN03). Briefly, mouse L cells transfected using polybrene with a lentivirus containing the *homo sapiens CD276* transcript variant 1 mRNA (NM_001024736.1) (Genecopia LPP-Z3060-LV105-100) and selected using puromycin suppression, were used to immunize New Zealand White mice (NZBWF1/J, Jackson Laboratory) or Balb/c (Charles River) by foot-pad immunization ^129^. Sera from immunized mice were screened by ELISA for differential binding to human 4Ig-B7-H3-expressing L cells compared to both L cells transfected with the vector control transcript (Genecopia LPP-NEG-Lv105-025-c) and parental L cells as well as binding to mouse 2Ig-B7-H3 protein (R&D 1397-B3-050). Selected clones were used to generate stable hybridomas. Monoclonal antibodies obtained from the supernatants of these hybridomas were further evaluated for human, mouse and porcine 4Ig-/2Ig-B7-H3 affinity and specificity.

MIL33B was produced by purification of supernatants from the MIL33B hybridoma by either the MDACC Monoclonal Antibody Core or by BioXCell. The affinity of each MIL33B stock for human and mouse 4Ig-/2Ig-B7-H3 was validated by ELISA.

### ELISA and Bi-Layer Interferometry Analysis

ELISA was performed using MIL33B to probe plate-bound human and mouse B7-H3 extracellular domains (human 4Ig-B7-H3 protein (R&D 2318-B3-050); mouse 2Ig-B7-H3 protein (R&D 1397-B3-050)), with 1:2000 dilutions of secondary antibody, (goat anti-mouse IgG (Fc) HRP (Biorad)). To further characterize antibodies, binding kinetics determination was performed using the OCTET RED384 platform, a technology based on Bio-Layer Interferometry (BLI). Briefly, the mouse anti-B7-H3 clone, MIL33B, was immobilized on biosensors (AMC) using anti-mouse IgG Fc capture at a fixed concentration of 10 μg/ml. Loaded biosensors then interacted with 7 serial dilutions (ranging from 3 nM to 200 nM) of the extracellular domain of various B7-H3 isoforms or relevant B7 protein homologues. The BLI system measured association and dissociation constants, Ka and Kd, respectively, and calculated affinity constants (K_D_) using the Data Acquisition software.

### Human Tissue Microarray Fluorescence Staining

Frozen human tissue microarrays were purchased from Fisher Scientific (#50-180-886) containing arrayed normal and cancer tissues for multiple organ sites. Staining was performed following a brief fix (5 minutes) at room temperature of the slide in 4% paraformaldehyde. Following fixation, slides were washed two times in 5%BSA/PBS for 5 minutes each. Blocking was performed in 5% BSA/PBS containing 1:100 IgG2a (Final concentration∼ 50 µg/mL) at room temperature for 30 minutes. Primary antibody was added in 1% BSA/PBS and incubated overnight at 4°C protected from light. MIL33B-AF594 and IgG2a-AF594 were diluted 1:100; final concentration was 10 µg/mL. Following overnight incubation, slides were washed 2X in 1%BSA/PBS for 5 minutes each. Nuclear staining was performed with DAPI at a dilution of 1:1000 in 1%BSA/PBS for 10 minutes at room temperature. Slides were washed an additional time in 1%BSA/PBS and mounted using antifade mounting media. Once dried, the slides were sealed and images were captured on a Nikon TiE Epifluorescence microscope.

### Cell Line Generation

B16F10 murine melanoma cells, 4T1 murine triple negative breast cancer cells, and CT26 murine colorectal carcinoma cells were transfected using polybrene with lentiviral constructs containing the *homo sapien CD276* transcript variant 1 mRNA (NM_001024736.1) (Genecopia LPP-Z3060-LV105-100) or the negative vector control transcript (Genecopia LPP-NEG-Lv105-025-c) and selected in puromycin. CD276^−/−^ cells were generated by transfecting HeLa parental cells with human-derived *CD276* Crispr/Cas9 pooled plasmids (Santa Cruz Biotechnology sc-402032) or double nickase control plasmids (each containing a GFP marker, three separate guide RNAs, and a plasmid encoding Cas9). GPF-positive cells were sorted and expanded. HCT116 cells or HCT116 cells stably transfected with a *KB5-IkBα-FLUC* reporter ^130^ were cultured as described.

### Live Cell Fluorescence Microscopy

Cells were plated in a 4-well Ibidi dish at 5×10^4^ cells in 500 µL of media. Cells were incubated at 37 °C in 5% CO_2_ for 24-48 hours. MIL33B or mouse isotype control IgG2a (BioXcell BE0085) were labeled with Alexa594 (Thermofischer A20185) by the MDACC flow cytometry core. Cells were subsequently incubated with 10 µg/mL of either Alexa549-labeled MIL33B or IgG2a and 5 µL of a 1:100 dilution in PBS of Hoechst stain for 1 hour and imaged on a Nikon TiE inverted microscope with a 40x objective.

### Antibody Conjugation with p-SCN-Bn-Deferoxamine or p-SCN-Bn-DOTA

For antibodies conjugated with p-SCN-Bn-Deferoxamine (Macrocyclics), 4-7mg/mL of either MIL33B or mouse IgG2a (BioXcell) were desalted into 0.1M NaHCO_3_ (pH = 8.4) using 7K MWCO Zeba (Thermofisher) protein desalting spin columns. When conjugating MIL33B to p-SCN-Bn-DOTA (Macrocyclics), the stock solution was concentrated to 9-11 mg /mL using Amicon Ultra-0.5 centrifugal filter units with a 30 kDa cutoff, and then similarly desalted into a 0.1 M solution of sodium bicarbonate (pH = 8.4). Antibodies were incubated with 5 meq of each respective chelator dissolved in DMSO for 1 hour at 37°C with periodic mixing. Subsequently, antibodies were desalted into 0.1 M ammonium acetate (pH = 6.6-7) as described previously ^131^.

### Radiolabeling of MIL33B with Zirconium-89

Zirconium-89 in 1 M oxalic acid was produced by the MDACC cyclotron facility or the University of Wisconsin cyclotron facility. Zirconium-89 oxalate (3 mCi) was neutralized to pH 7 by 1.0 M Na_2_CO_3_ (pH = 11). Following, 20-30 µL of a 4-7 mg/mL solution of DFO-conjugated antibodies, MIL33B or mouse IgG2a, was added to the buffer and the final volume brought to 200 µL with PBS and with zirconium-89 oxalate for 1 hour at 37 °C. Chelation efficiency was evaluated by radio-TLC with antibody solutions quenched in 50 mM DPTA (pH = 7) using 50 mM DPTA (pH = 7) as the running solvent. Chelated antibodies were purified into PBS using a PD-10 column.

### Radiolabeling of MIL33B with Yttrium-90

[^90^Y]YCl_3_ in 0.04M HCl was purchased from Eckert & Ziegler Radiopharma. [^90^Y]Yttrium chloride was buffered in an equal volume of 0.1M ammonium acetate (pH = 5.6). Following, 50 µL of DOTA-MIL33B was added to the solution. DOTA-MIL33B was incubated with [^90^Y]yttrium chloride for 1 hour at 37°C. Chelation efficiency was evaluated by radio-TLC with antibody solutions quenched in 50 mM EDTA using 10 mM EDTA as the running solvent. ^90^Y-DOTA-MIL33B were purified into PBS using a PD-10 column.

### Tumor Models

All experiments were approved under an MDACC IACUC protocol 00001179-RN01 and RN02. Mice aged 5-12 weeks were used for all experiments. Tumor cells were implanted subcutaneously on the flanks of female Athymic Nude Mice 490 immunodeficient mice (Charlies River) or flanks of respective immunocompetent female Balb/c mice (Taconic) or female C57BL/6 mice (Taconic). For all mice, tumors were measured using calipers twice weekly. Mice were defined to reach endpoint when either the length or width of the tumor reached 15 mm or the tumors had ulcers of 3 mm, at which point mice were euthanized by CO_2_ asphyxiation and cervical dislocation.

### PET-CT Imaging Methods and Analysis

Mice were injected with approximately 20-60 μCi of either ^89^Zr-DFO-MIL33B or ^89^Zr-DFO-IgG2a intravenously. For cold blocking experiments, mice received 200 µg of un-labeled MIL33B 1 hour before injection with the tracer of interest. Mice were imaged at 24 hours, 72 hours and 144 hours on a PET-CT scanner (Albira, Bruker), with a field of view of 120 mm for PET and 70-75 mm for CT. Mice were imaged with a single 10 minute PET scan and a fast CT HV, HD scan. VOIs were drawn in PMOD and used to calculate %ID/cc for respective VOIs at each time point.

### Cherenkov Radiation Optical Imaging

Images of mice were obtained using an IVIS Spectrum, open filter, FOV 23 cm x 23 cm, and image acquisition time of 5 minutes, followed by acquisition of a bright light image.

### ^90^Y-DOTA-MIL33B Treatment Strategy

Mice harboring CT26 4Ig-B7-H3 or neg vector tumors were selected for tumors with a volume of about 50 mm^3^ - 150 mm^3^ at 10-12 days post tumor implantation and randomized into treatment groups. Untreated mice were followed regardless of initial tumor size. Mice were treated with intravenous injection of 100 µCi of ^90^Y-DOTA-ML33B or 100 µL of sterile saline. For HeLa B7-H3^+/+^ WT or HeLa B7-H3^−/−^ KO, 5×10^6^ cells were implanted subcutaneously in the flanks of athymic nude mice. Mice harboring HeLa B7-H3^+/+^ or HeLa B7-H3^−/−^ KO tumors were not size selected and received an intravenous injection of 100 µCi of ^90^Y-DOTA-ML33B 30 days post tumor implantation or were not treated (serving as a tumor growth rate control), thus anchoring survival with previously performed animal studies ^52^.

### Tumor Growth Rate Comparison

In comparing the growth rates of HeLa B7-H3^+/+^ WT or HeLa B7-H3^−/−^ KO tumors treated with ^90^Y-DOTA-ML33B i.v., changes in tumor growth were calculated on a mouse by mouse basis by calculating the change in tumor size at each time point compared to initial tumor size **(Supplemental Figure 9)**. Slopes of tumor growth (growth rates) were calculated with a 3-point exponential fitting curve using Prism software.

### Immune Cell Depletion In Vivo

Mice harboring CT26 4Ig-B7-H3 tumors were treated with a single dose of 100 µCi of ^90^Y-DOTA-MIL33B i.v. 1 day before treatment, mice received either 250 µg of anti-mouse CD8b (clone 53-5.8 BioXcell), anti-mouse CD4 (clone GK1.5 BioXcell), or anti-rat IgG polyclonal (BioXcell) i.p. and then received subsequent doses of each respective depleting antibody 2 days, 6 days and 9 days post treatment with ^90^Y-DOTA-MIL33B i.v. Response to treatment with depleting antibodies was monitored by caliper measurements of the tumors.

### Tumor Cell Re-Challenge

Mice harboring CT26 4Ig-B7-H3 tumors that had previously been treated with 100 µCi of ^90^Y-DOTA-MIL33B were followed for 100 days post tumor implantation. Mice whose tumors had regressed and did not recur at the 100-day time point were deemed long-term survivors. Long-term survivors were implanted with 5×10^4^ CT26 4Ig-B7-H3 cells in 100 µL of a 1:1 ratio of PBS:matrigel subcutaneously on the left flank. Naïve female Balb/c mice (Taconic Biosciences) were also implanted identically with 5×10^4^ CT26 4Ig-B7-H3 cells in 100 µL of a 1:1 ratio of PBS:matrigel subcutaneously on the left flank. Response to the tumor cell re-challenge was monitored by caliper measurements.

### MIL33B Monotherapy

Balb/c female mice (Taconic biosciences) were implanted with 1×10^5^ CT26 4Ig-B7-H3 cells in 100 µL PBS subcutaneously in the right flank. Mice received 200 µg i.p. of either MIL33B, mouse IgG2a isotype control antibody (Bioxcell), or 100 µL PBS on day 3, 6, 9, 12, 15, 18, 21, and 24 post cell implantation.

### Histology

Mice harboring CT26 4Ig-B7-H3 tumors were treated with 100 µCi of ^90^Y-DOTA-MIL33B i.v. according to the protocol. 6 days after treatment, mice were sacrificed, tumors fixed in formalin for 24-36 hours and then subsequently stored in 70% ethanol for radioactive decay. Tumors were stained by H&E staining, anti-CD4 (Abcam ab183685), anti-CD3 (Cell Signaling 99940), anti-CD8 (Abcam ab217344), anti-Ki67 (Cell Signaling 12202), anti-CD31 (abcam ab124432) and anti-cleaved caspase-3 (Cell Signaling 9662) by the MDACC histology core facility. Histopathology review, including immuno-histochemistry scoring, was performed by a board-certified pathologist.

### Statistics

Statistics were calculated using GraphPad Prism 9.0.0 (121). Data are represented as mean ± standard error. Pairs were compared using a student’s t-test where p-values less than or equal to 0.05 were considered significant.

### Data Availability

All source data is contained within the text and supplemental files or tables.

## Acknowledgements

Sarah Glazer was supported by an NCI pre-doctoral fellowship (5F30CA239332). Margie Sutton was supported by an NCI post-doctoral fellowship (F32CA250323). The MDACC Small Animal Imaging Facility, Flow Cytometry and Cellular Imaging Core, Research Histology Core Laboratory, and Monoclonal Antibody Facility are supported by a Cancer Center Support Grant (P30 CA016672). We would like to thank members of the Small Animal Imaging Facility (Charles Kingsley, Jorge Delacerda, Emily Newson, and Jennifer Meyer) for their expert assistance with PET-CT imaging, and members of the Monoclonal Antibody Facility (Laura Bover, Felipe Amazy-Manazanares, and Long Vien) for their expert help in developing and producing anti-B7-H3 antibodies.

## Author Contributions

S.T.G. and D.P.W. designed the antibody immunization and screening strategy. S.E.G, S.T.G and D.P.W designed the *in vitro* and *in cellulo* selection strategy. S.E.G, F.P., S.T.G., and D.P.W. designed the radiolabeling and radiotherapy strategies. S.E.G, M.N.S., P.Y., F.P. and S.T.G. designed and executed experiments. All authors provided data analysis. S.E.G, S.T.G., M.N.S, and D.P.W. wrote the manuscript and all authors edited revised manuscripts. D.P.W. provided resources and reagents.

