## Supplemental Information for "Anti-Cancer Immune Priming with Beta-Radioligand Therapy and Isoform-Selective Targeting of 4Ig-B7-H3"

### Detailed Methods

*Antibody Development and MIL33B Production*

Anti-B7-H3 antibodies were generated at the MDACC Monoclonal Antibody Core Facility (protocol 00000620-RN01 through RN03). Briefly, mouse L cells transfected using polybrene with a lentivirus containing the homo *sapiens CD276* transcript variant 1 mRNA (NM_001024736.1) (Genecopia LPP-Z3060-LV105-100) and selected using puromycin suppression, were used to immunize New Zealand White mice (NZBWF1/J, Jackson Laboratory) or Balb/c (Charles River) by foot-pad immunization[119]. Sera from immunized mice were screened by ELISA for differential binding to human 4Ig-B7-H3-expressing L cells compared to both L cells transfected with the vector control transcript (Genecopia LPP-NEG-Lv105-025-c) and parental L cells as well as binding to mouse 2Ig-B7-H3 protein (R&D 1397-B3-050). Selected clones were used to generate stable hybridomas. Monoclonal antibodies obtained from the supernatants of these hybridomas were further evaluated for mouse and human 4Ig-/2Ig-B7-H3 affinity and specificity.

MIL33B was produced by purification of supernatants from the MIL33B hybridoma by either the MDACC Monoclonal Antibody Core or by BioXCell. The affinity of each MIL33B stock for human and mouse 4Ig-/2Ig-B7-H3 was validated by ELISA using MIL33B to probe plate-bound human and mouse B7-H3 extracellular domains (human 4Ig-B7-H3 protein (R&D 2318-B3-050); mouse 2Ig-B7-H3 protein (R&D 1397-B3-050)).

*ELISA*

For proteins, 96-well Costar plates were coated with 50 µL of 0.1 µg/mL of respective protein and incubated at room temperature overnight. For cells, $15x10^6$ cells were suspended in 4 mL of PBS containing 0.01% of $MgCl_2$ and $CaCl_2$. 40 µL of suspended cells were plated in a 96-well RIA plate (Costar 3369-Fisher) and dried overnight. Plates were stored at -20°C for at least 24 hours. Dried cells were rehydrated with 200 µL of PBS for 10 min at room temperature before starting the ELISA protocol. Wells were blocked with 200 µL of 2% PBSTB (0.05% Tween-20 and 2% BSA in PBS) at room temperature for 1 hour. Plates were washed three times with PBST (0.05% Tween-20 in PBS) and 50 µL per well of a dilution series of the antibody of interest were added to the plates. Plates were incubated at room temperature for 1 hour and then washed three times with PBST. 50 µL of 1:2000 dilutions of secondary antibody, (goat anti-mouse IgG (Fc) HRP (Biorad)), in PBSTB was added to each well and incubated at room temperature for 1 hour. Plates were then washed four times with PBS and 50 µL per well of TMB substrate was added to each well for 20-30 minutes at room temperature. Next, 50 µL per well of stop solution (0.2 M $H_2SO_4$) was added to each well. Plates were imaged on a Synergy H4 Hybrid Microplate Reader at 450 nm.

*Bi-Layer Interferometry: Octet Analysis*

To further characterize antibodies, binding kinetics determination was performed using the OCTET RED384 platform, a technology based on Bio-Layer Interferometry (BLI). Briefly, the mouse anti-B7-H3 clone, MIL33B, was immobilized on biosensors (AMC) using anti-mouse IgG Fc capture at a fixed concentration of 10 µg/ml. Loaded biosensors then interact with 7 serial dilutions (ranging from 3 nM to 200 nM) of the extracellular domain of various B7-H3 isoforms or relevant B7 protein homologues, human 2Ig-B7-H3 protein (R&D 1949-B3), mouse 2Ig-B7-H3 protein (R&D 1397-B3-050), human 4Ig-B7-H3 protein (R&D 2318-B3-050), mouse PD-L2 protein (R&D 9107-PL), human PD-L2 protein (R&D 9075-PL), human B7-H4 protein (R&D 6570-B7), mouse B7-H4 protein (R&D 2154-B7), human B7-H2 protein (R&D 8206-B7), mouse B7-H2 protein, (R&D 8127-B7), human PD-1 protein (R&D 8986-PD) and mouse PD-1 protein (R&D 9047-PD). Porcine B7-H3 protein was custom produced by Biotechne. The BLI system measured association and dissociation constants, Ka and Kd, respectively, and calculated affinity constants ($K_D$) using the Data Acquisition software.

*Cell Line Generation*

B16F10 murine melanoma cells, 4T1 murine triple negative breast cancer cells, and CT26 murine colorectal carcinoma cells were transfected using polybrene with lentiviral constructs containing the *homo sapien CD276* transcript variant 1 mRNA (NM_001024736.1) (Genecopia LPP-Z3060-LV105-100) or the negative vector control transcript (Genecopia LPP-NEG-Lv105-025-c) and selected in puromycin. Expression was confirmed by Western blot. Transfected B16F10 cells were cultured in 2 µg/mL puromycin in DMEM and 10% FBS. Transfected 4T1 cells were cultured in 3 µg/mL puromycin in RPM1 1640 with glutamine, sodium pyruvate and 10% FBS. Transfected CT26 cells were cultured in 6.4 µg/mL puromycin with MEM, sodium pyruvate, non-essential amino acids and 10% FBS. $CD276^{-/-}$ cells were generated by transfecting HeLa parental cells with human-derived *CD276* Crispr/Cas9 pooled plasmids (Santa Cruz Biotechnology sc-402032) or double nickase control plasmids (each containing a GFP marker, three separate guide RNAs, and a plasmid encoding Cas9). GPF-positive cells were sorted and expanded. Single cell clones were grown and screened for CD276 knockout or established from control cells. Cells were maintained in DMEM supplemented with 10% FBS and 1% L-Glutamine.

HCT116 cells or HCT116 cells stably transfected with a *KB5-IkBα-FLUC* reporter [120] were cultured in McCoy 5A supplemented with 10% FBS and 0.3 µg/mL puromycin.

*Western Blot*

Cells were plated in 100 mm cell culture dishes. 24 hours later, cells were washed two times with PBS and 500 µL of lysis buffer (MCLB buffer and a protease inhibitor) added directly to the dish. Cell fragments were scraped from the dish and placed on ice for 20 min. Subsequently, cells were centrifuged at 15,000 rpm at 4 °C for 20 min, and the supernatant was retrieved and stored at -80°C for downstream use.

20-40 µg of protein, determined by the Bradford assay, in a 1:1 ratio of Laemmli buffer with β-mercaptoethanol were denatured at 95 °C for 5 min. Samples or a PageRule Plus Ladder (Thermofisher 26619) were loaded in a protean TGx precast 4-20% polyacrylamide gel (Bio-Rad 4568095). Gels were run in a 1x Tris/Glycine/SDS buffer at 100-200 volts and transferred to a 0.2 mm PVDF membrane using the BioRad Turbo Transfer System. Membranes were blocked in blocking buffer (0.5 g nonfat milk powder in 10 mL TBST) for 1 hour washed three times in 1x TBST for 10 minutes each ,and incubated with 1 µg/mL of anti-B7-H3 antibody (AF1397 R&D Systems) or a 1:10000 dilution of anti-GAPDH (Abcam ab181602) in blocking buffer and incubated overnight. Membranes were washed three times with TBST and incubated with a 1:5000 dilution of anti-goat-HRP (Sigma A5240) or anti-Rabbit-HRP (BioRad) for 1 hour. Then washed three times with TBST, and then incubated with Pierce West Pico ECL for 5 minutes. Membranes were imaged on an Azure c600 Western blot imaging, chemiluminescence detection system.

*Human Tissue Microarray Fluorescence Staining*

Frozen human tissue microarrays were purchased from Fisher Scientific (#50-180-886) containing arrayed normal and cancer tissues for multiple organ sites. Staining was performed following a brief fix (5 minutes) at room temperature of the slide in 4% paraformaldehyde. Following fixation, slides were washed two times in 5%BSA/PBS for 5 minutes each. Blocking was performed in 5% BSA/PBS containing 1:100 IgG2a (Final concentration~ 50 µg/mL) at room temperature for 30 minutes. Primary antibody was added in 1% BSA/PBS and incubated overnight at 4ºC protected from light. MIL33B-AF594 and IgG2a-AF594 were diluted 1:100; final concentration was 10 µg/mL. Following overnight incubation, slides were washed 2X in 1%BSA/PBS for 5 minutes each. Nuclear staining was performed with DAPI at a dilution of 1:1000 in 1%BSA/PBS for 10 minutes at room temperature. Slides were washed an additional time in

1%BSA/PBS and mounted using antifade mounting media. Once dried, the slides were sealed and images were captured on a Nikon TiE Epifluorescence microscope.

*Live Cell Fluorescence Microscopy*

Cells were plated in a 4-well Ibidi dish at $5x10^4$ cells in 500 µL of media. Cells were incubated at 37 °C in 5% $CO_2$ for 24-48 hours. MIL33B or mouse isotype control IgG2a (BioXcell BE0085) were labeled with Alexa594 (Thermofischer A20185) by the MDACC flow cytometry core. Cells were subsequently incubated with 10 µg/mL of either Alexa549 labeled MIL33B or IgG2a and 5 µL of a 1:100 dilution in PBS of Hoechst stain for 1 hour. Cells were washed two times with PBS and placed in colorless media with 10% FBS. Cells were imaged on a Nikon TiE inverted microscope with a 40x objective with the following acquisition rates: 100 ms DIC, 100 ms DAPI and 1 sec TRITC. For blocking studies, cells were pre-incubated with 50 µg/mL of MIL33B or mouse IgG2a isotype control antibody for 40 minutes at 37°C in 5% CO2 and not washed before incubating with the relevant Alexa549-labeled antibodies.

*Antibody Conjugation with p-SCN-Bn-Deferoxamine or p-SCN-Bn-DOTA*

For antibodies conjugated with p-SCN-Bn-Deferoxamine (Macrocyclics), 4-7mg/mL of either MIL33B or mouse IgG2a (BioXcell) were desalted into 0.1M $NaHCO_3$ (pH = 8.4) using 7K MWCO Zeba (Thermofisher) protein desalting spin columns. When conjugating MIL33B to p-SCN-Bn-DOTA (Macrocyclics), the stock solution was concentrated to 9-11 mg /mL using Amicon Ultra-0.5 centrifugal filter units with a 30 kDa cutoff, and then similarly desalted into a 0.1 M solution of sodium bicarbonate (pH = 8.4). Antibodies were incubated with 5 meq of each respective chelator dissolved in DMSO for 1 hour at 37°C with periodic mixing. Subsequently, antibodies were desalted into 0.1 M ammonium acetate (pH = 6.6-7) as described previously [121]. Final conjugated antibody concentrations were determined by spectrophotometry (NanoDrop) and antibodies were stored at 4°C for downstream experiments.

*Radiolabeling of MIL33B with Zirconium-89*

Zirconium-89 in 1 M oxalic acid was produced by the MDACC cyclotron facility or the University of Wisconsin cyclotron facility. Zirconium-89 oxalate (3 mCi) was neutralized to pH 7 by 1.0 M $Na_2CO_3$ (pH = 11). Following, 20-30 µL of a 4-7 mg/mL solution of DFO-conjugated antibodies, MIL33B or mouse IgG2a, was added to the buffer and the final volume brought to 200 µL with PBS and with zirconium-89 oxalate for 1 hour at 37 °C. Chelation efficiency was evaluated by radio-TLC with antibody solutions quenched in 50 mM DPTA (pH = 7) using 50 mM DPTA (pH = 7) as the running solvent. Chelated antibodies were purified into PBS using a PD-10 column. Purity was validated by radio-TLC and radio-SEC-HPLC with a 200mm 10-30 SuperDex column using a 71 minute isocratic elution in PBS with 0.1M NaCl and 0.05% sodium azide. Specific activity was determined by comparing the area under the curve of the 280 nm UV spectrum of the antibody obtained from radio-SEC-HPLC to a standard curve of MIL33B obtained from a dose dilution of the antibody.

*Radiolabeling of MIL33B with Yttrium-90*

[$^{90}$Y]$YCl_3$ in 0.04M HCl was purchased from Eckert & Ziegler Radiopharma. [$^{90}$Y]Yttrium chloride was buffered in an equal volume of 0.1M ammonium acetate (pH = 5.6). Following, 50 µL of DOTA-MIL33B was added to the solution. DOTA-MIL33B was incubated with [$^{90}$Y]yttrium chloride for 1 hour at 37°C. Chelation efficiency was evaluated by radio-TLC with antibody solutions quenched in 50 mM EDTA using 10 mM EDTA as the running solvent. $^{90}$Y-DOTA-MIL33B were purified into PBS using a PD-10 column. Purity was validated by radio-TLC and radio-SEC-HPLC with a 200mm 10-30 SuperDex column using a 71 minute isocratic elution in PBS with 0.1M NaCl and 0.05% sodium azide. De-chelation of $^{90}$Y-DOTA-MIL33B was evaluated immediately after radiolabeling by radio-SEC-HPLC. Specific activity was determined by comparing the area under the curve of the 280 nm UV spectrum of $^{90}$Y-DOTA-MIL33B obtained from radio-SEC-HPLC to a standard curve of MIL33B obtained from a dose dilution of the antibody.

*Tumor Models*

All experiments were approved under an MDACC IACUC protocol 00001179-RN01 and RN02. Mice aged 5-12 weeks were used for all experiments. For experiments using HeLa B7-H3$^{+/+}$ WT or HeLa B7-H3$^{-/-}$ KO tumors, 5x10$^6$ cells in 100 µL PBS of either HeLa B7-H3$^{+/+}$ cells or HeLa B7-H3$^{-/-}$ cells were implanted subcutaneously on the right flank of female Athymic Nude Mice 490 Homozygous, Immunodeficient (Charlies River). For experiments using 4T1 4Ig-B7-H3 tumors or 4T1 neg vector tumors, 1x10$^4$ cells in 100 µL of PBS of 4T1 4Ig-B7-H3 cells were implanted subcutaneously on the right flank and 4T1 neg vector cells were implanted subcutaneously on the left flank of female Balb/c mice (Taconic). For experiments using B16F10 4Ig-B7-H3 or neg vector tumors, 1x10$^4$ cells in 100 µL PBS of either B16F10 4Ig-B7-H3 cells or B16F10 neg vector cells were implanted subcutaneously on the right flank of female C57BL/6 mice (Taconic). For experiments using CT26 4Ig-B7-H3 or neg vector tumors, 1-5x10$^4$ cells in 100 µL of a 1:1 solution of PBS and matrigel or 1e5 cells in 100 µL PBS for mice treated with MIL33B monotherapy were implanted subcutaneously on the right flank of female Balb/c mice (Taconic). For all mice, tumors were measured using calipers twice weekly. Tumor volume was calculated by the equation, $(l \times w^2)/2$, where l is the length and w is the width of the tumor. Mice were defined to reach endpoint when either the length or width of the tumor reached 15 mm or the tumors had ulcers of 3 mm, at which point mice were euthanized by $CO_2$ asphyxiation and cervical dislocation.

*PET/CT Imaging Methods and Analysis*

Mice were injected with approximately 20-60 µCi of either $^{89}$Zr-DFO-MIL33B or $^{89}$Zr-DFO-IgG2a intravenously. Injected amounts were determined by measuring the amount of tracer in the syringe before and after injection. For cold blocking experiments, mice received 200 µg of un-labeled MIL33B 1 hour before injection with the tracer of interest. Mice were imaged at 24 hours, 72 hours and 144 hours on a PET/CT scanner (Albira, Bruker), with a field of view of 120 mm for PET and 70-75 mm for CT. Mice were imaged with a single 10 minute PET scan and a fast CT HV, HD scan. VOIs were drawn in PMOD and used to calculate %ID/cc for respective VOIs at each time point.

*Cherenkov Radiation Optical Imaging*

Mice injected with 100 µCi $^{90}$Y-DOTA-ML33B i.v. were anesthetized with 2% isoflurane, and imaged on an IVIS Spectrum at the following timepoints post injection: 2, 7, 10, 14, 17, and 24 days. Images were obtained using an open filter, FOV 23 cm x 23 cm, and image acquisition time of 5 minutes, followed by acquisition of a bright field image.

*$^{90}$Y-DOTA-MIL33B Treatment Strategy*

Mice harboring CT26 4Ig-B7-H3 or neg vector tumors were selected for tumors with a volume of about 50 mm$^3$ - 150 mm$^3$ at 10-12 days post tumor implantation and randomized into treatment groups. Untreated mice were followed regardless of initial tumor size. Mice were treated with intravenous injection of 100 µCi of $^{90}$Y-DOTA-ML33B or 100 µL of sterile saline. Mice harboring HeLa B7-H3$^{+/+}$ or HeLa B7-H3$^{-/-}$ KO tumors were not size selected and received and intravenous injection of 100 µCi of $^{90}$Y-DOTA-ML33B 30 days post tumor implantation. Final tracer dosing was determined by measuring the amount of tracer in the syringe before and after injection.

*Tumor Growth Rate Comparison*

In comparing the growth rates of HeLa B7-H3$^{+/+}$ WT or HeLa B7-H3$^{-/-}$ KO tumors treated with $^{90}$Y-DOTA-ML33B i.v., changes in tumor growth was calculated on a mouse by mouse basis by calculating the change in tumor size at each time point compared to initial tumor size. Slopes of tumor growth, or growth rates, were calculated with a 3-point exponential fitting cure in Prism, one KO mouse was excluded from the analysis due to lack of sufficient points in the curve for a curve fit in the initial slope regime. Exponential slopes in each group were compared using a 2-way ANOVA 15 days before treatment and 15 days after treatment.

*Cell Depletion In Vivo*

Mice harboring CT26 4Ig-B7-H3 tumors were treated with a single dose 100 µCi of $^{90}$Y-DOTA-MIL33B i.v. 1 day before treatment, mice received either 250 µg of anti-mouse CD8b (clone 53-5.8 BioXcell), anti-mouse CD4 (clone GK1.5 BioXcell), or anti-rat IgG polyclonal (BioXcell) i.p. and then received subsequent doses of each respective depleting antibody 2 days, 6 days and 9 days post treatment with $^{90}$Y-DOTA-MIL33B i.v. Response to treatment with depleting antibodies was monitored by caliper measurements of the tumors.

*Tumor Cell Re-Challenge*

Mice harboring CT26 4Ig-B7-H3 tumors that had previously been treated with 100 µCi of $^{90}$Y-DOTA-MIL33B were followed for 100 days post tumor implantation. Mice whose tumors had regressed and did not recur at the 100-day time point were deemed long-term survivors. Long-term survivors were implanted with $5x10^4$ CT26 4Ig-B7-H3 cells in 100 µL of a 1:1 ratio of PBS:matrigel subcutaneously on the left flank. Naïve female Balb/c mice (Taconic Biosciences) were also implanted identically with $5x10^4$ CT26 4Ig-B7-H3 cells in 100 µL of a 1:1 ratio of PBS:matrigel subcutaneously on the left flank. Response to the tumor cell re-challenge was monitored by caliper measurements.

*MIL33B Monotherapy*

Balb/c female mice (Taconic biosciences) were implanted with $1x10^5$ CT26 4Ig-B7-H3 cells in 100 µL PBS subcutaneously in the right flank. Mice received 200 µg i.p. of either MIL33B, mouse IgG2a isotype control antibody (Bioxcell), or 100 µL PBS at 3, 6, 9, 12, 15, 18, 21, and 24 days post cell implantation.

*Histology*

Mice harboring CT26 4Ig-B7-H3 tumors were treated with 100 µCi of $^{90}$Y-DOTA-MIL33B i.v. according to the protocol. 6 days after treatment, mice were sacrificed and tumors fixed in formalin for 24-36 hours and then subsequently stored in 70% ethanol for radioactive decay. Tumors were stained by H&E staining, anti-CD4 (Abcam ab183685), anti-CD3 (Cell Signaling 99940), anti-CD8 (Abcam ab217344), anti-Ki67 (Cell Signaling 12202), anti-CD31 (abcam ab124432) and anti-cleaved caspase-3 (Cell Signaling 9662) by the MDACC histology core facility. Histopathology review, including immuno-histochemistry scoring, was performed by a board-certified pathologist. Scores were then correlated on a per animal basis with rank order correlation.

*Statistics*

Statistics were calculated using GraphPad Prism 9.0.0 (121). Data are represented as mean $\pm$ standard error. Pairs were compared using a student's t-test where p-values less than or equal to 0.05 were considered significant.

*Figures*

All diagrams were created with Biorender.com software.

**Supplemental Figures:**

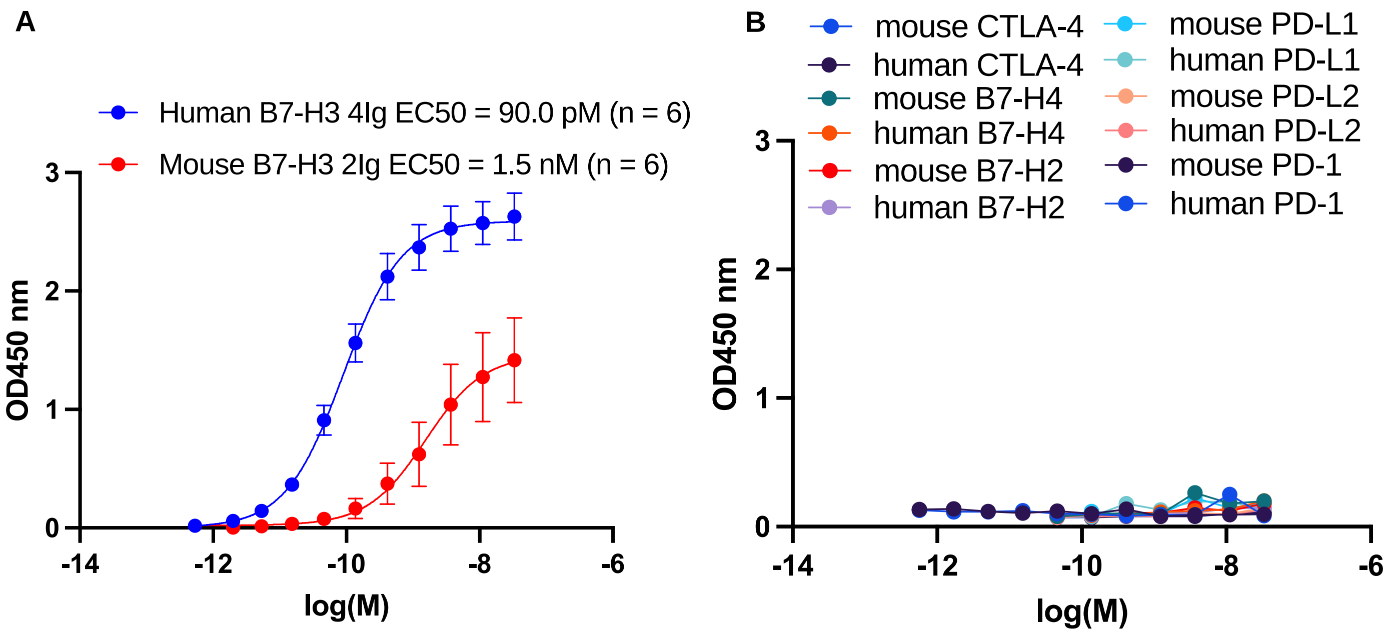


**Supplemental Figure 1: MIL33B demonstrated high affinity binding to human 4Ig-B7-H3 and lower affinity to mouse and human B7 homologues and relevant checkpoint proteins.** (A) ELISA of serial dilutions of MIL33B from different production lots demonstrated high affinity to plate-bound extracellular domain of human 4Ig-B7-H3 (EC50 = 90.0 pM, 95% CI: 62 pM – 130 pM; n = 6), and lower affinity to plate-bound extracellular domain of mouse 2Ig-B7-H3 (EC50 = 1.7nM, 95% CI: 0.47 nM – 5.0 nM; n = 6). (B) ELISA of MIL33B to plate-bound extracellular domains of mouse and human B7 family homologues and common immune checkpoint proteins demonstrated no substantial affinity (n = 1 for each homologue).

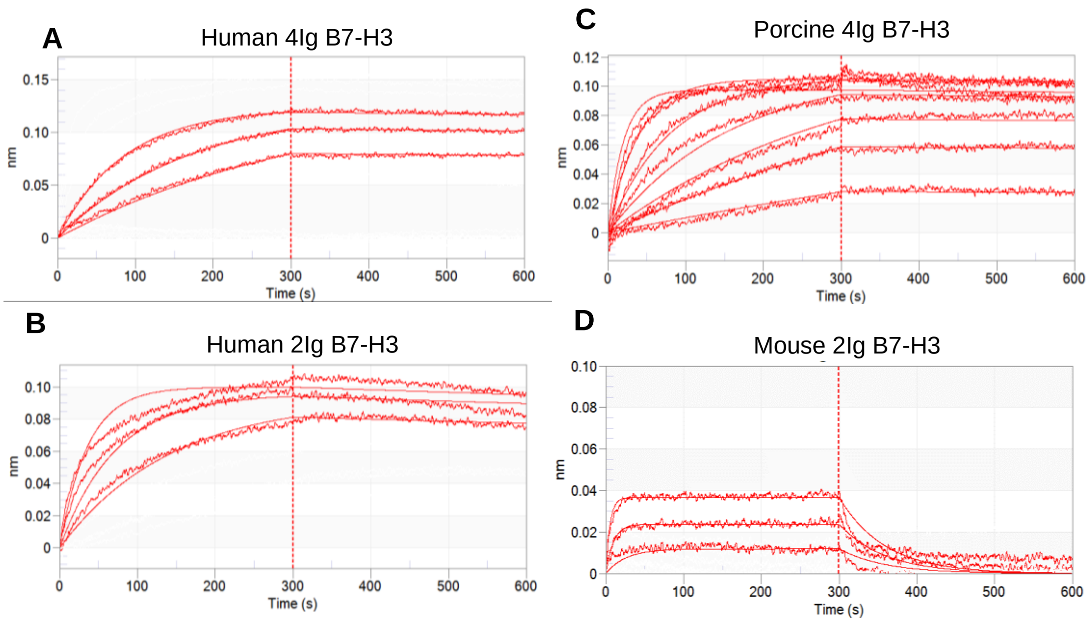


**Supplemental Figure 2: Biolayer interferometry of MIL33B binding to human, mouse and porcine B7-H3 isoforms.** Biolayer interferometry analysis of MIL33B binding to human 4Ig-B7-H3 (A), human 2Ig-B7-H3 (B), porcine 4Ig-B7-H3 (C), and mouse 2Ig-B7-H3 (D) demonstrated picomolar affinity for human 4Ig-B7-H3 and nanomolar affinity for mouse 2Ig-B7-H3.

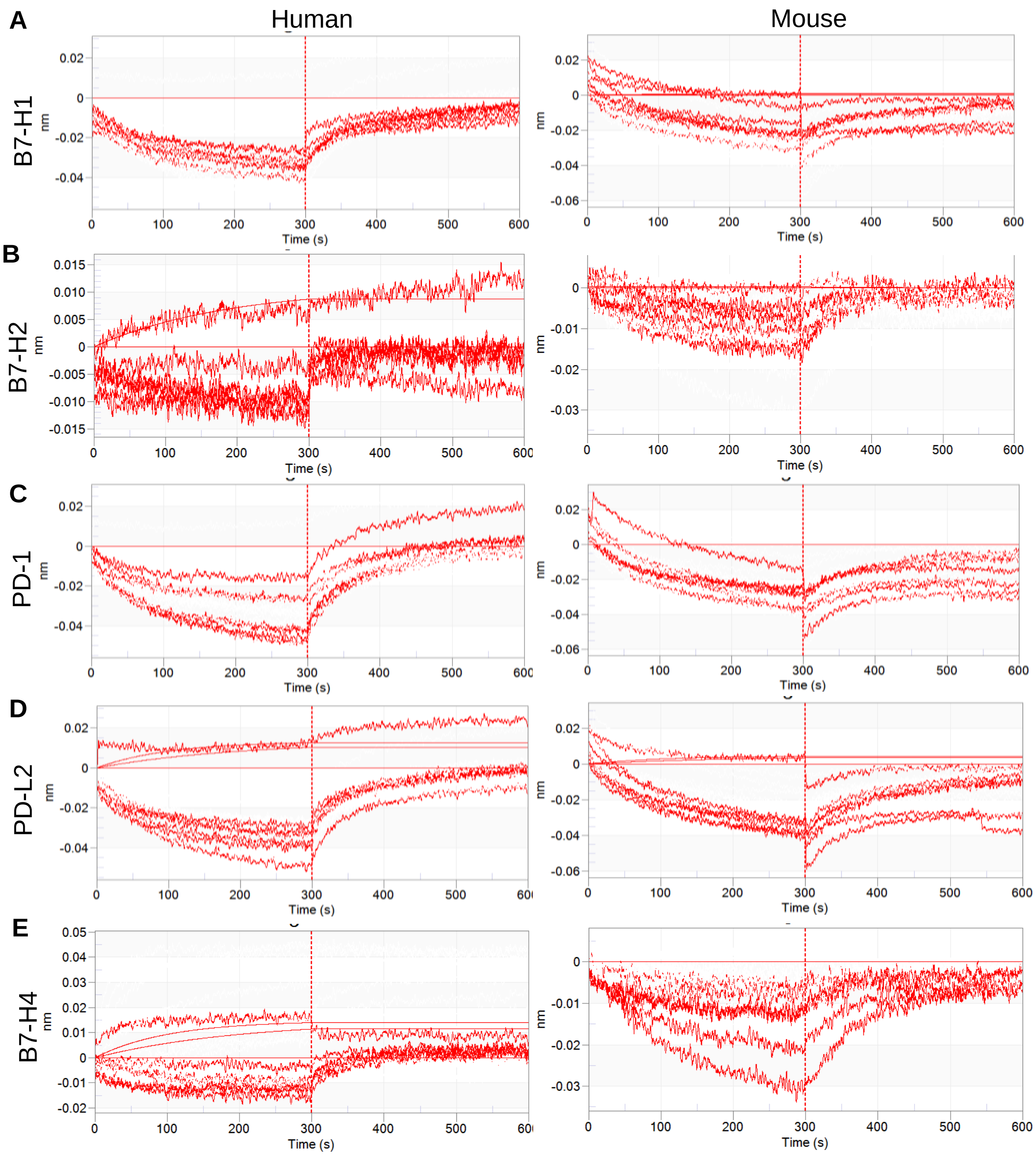


**Supplemental Figure 3: Biolayer interferometry of MIL33B binding to mouse and human B7 homologues and immune checkpoint proteins**. Curve fitting of MIL33B to immobilized human and mouse extracellular domains of B7-H1 (A), B7-H2 (B), PD-1 (C), PD-L1 (D) PD-L2 and B7-H4 (E), demonstrating no substantial binding.

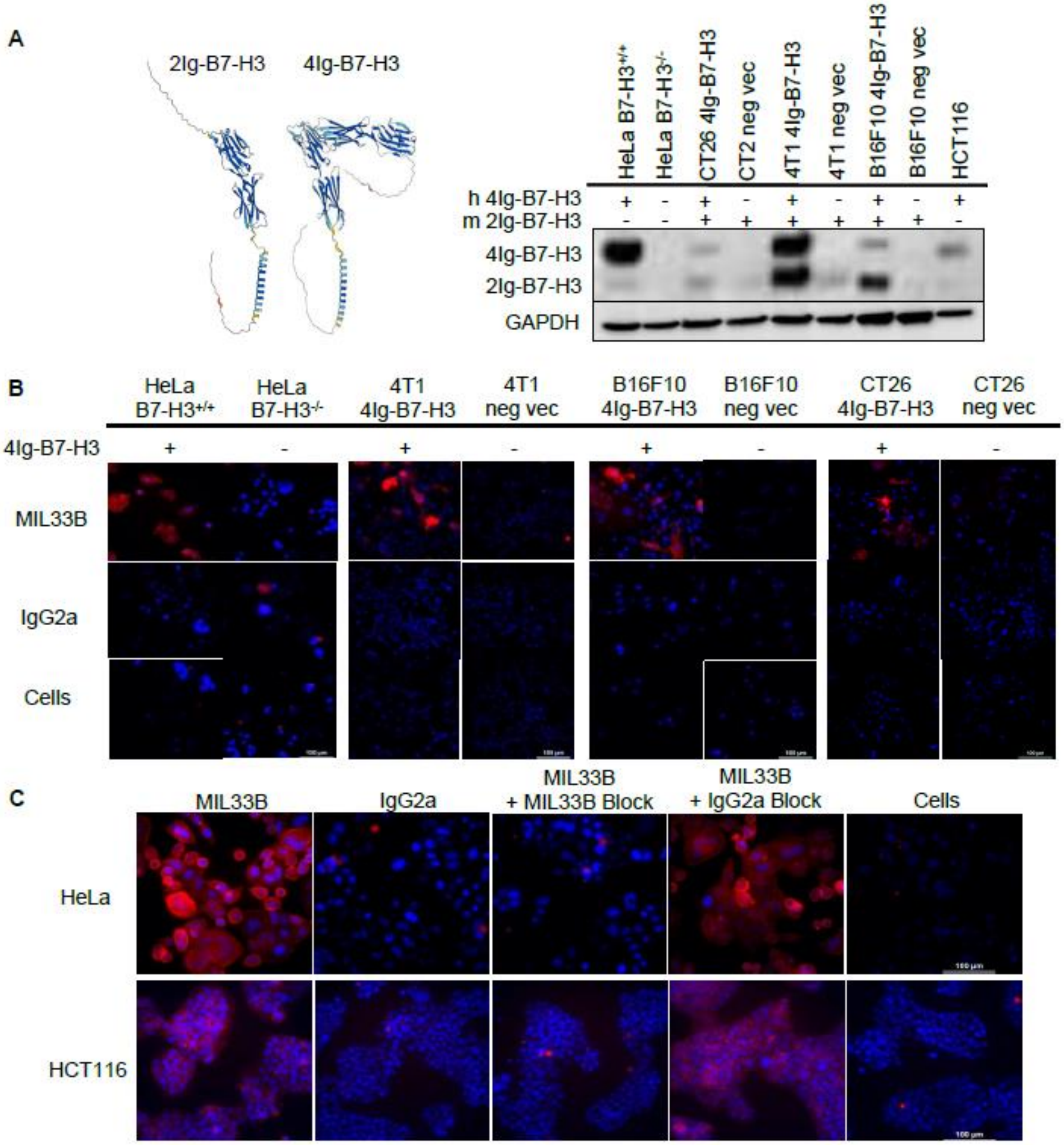


**Supplemental Figure 4. MIL33B has high affinity and specificity for the human 4Ig-B7-H3 isoform.** (**A**) Structure prediction (Alphafold V2) of mouse (m) 2Ig-B7-H3 and human (h) 4Ig-B7-H3 (left). Western blot for B7-H3 demonstrates a decrease in both 4Ig and 2Ig isoforms of B7-H3 in HeLa B7-H3-/- KO cells compared to HeLa B7-H3+/+ cells and an increase in human 4Ig-B7-H3 expression in transduced CT26 4Ig-B7-H3, 4T1 4Ig-B7-H3 and B16F10 4Ig-B7-H3 cells compared to cells transfected with the negative vector control: CT26 neg vec, 4T1 neg vec and B16F10 neg vec (right). (**B**) Alexa549-labeled MIL33B was incubated with human and murine cell lines in which human B7-H3 had been knocked out or in which cells had been transduced to express human 4Ig-B7-H3. MIL33B-Alexa549 demonstrated high cell-specific binding to HeLa B7-H3+/+ cells, 4T1 4Ig-B7-H3 cells, B16F10 4Ig-7B-H3, and CT26 4Ig-B7-H3 cells compared to HeLa-/- cells, 4T1 neg vec cells, B16F10 neg vec cells and CT26 neg vec cells or compared to binding of Alexa549-labeled mouse isotype control (IgG2a), or cells alone. (**C**) High 4Ig-B7-H3-expressing human HeLa and HCT116 cells incubated with Alexa549-labeled MIL33B demonstrated high intensity membrane-specific localization compared to Alexa549-labeled IgG2a or cells alone. Pre-incubation with un-labeled MIL33B was superior to murine IgG2a in displacing Alexa549-labeled MIL33B.

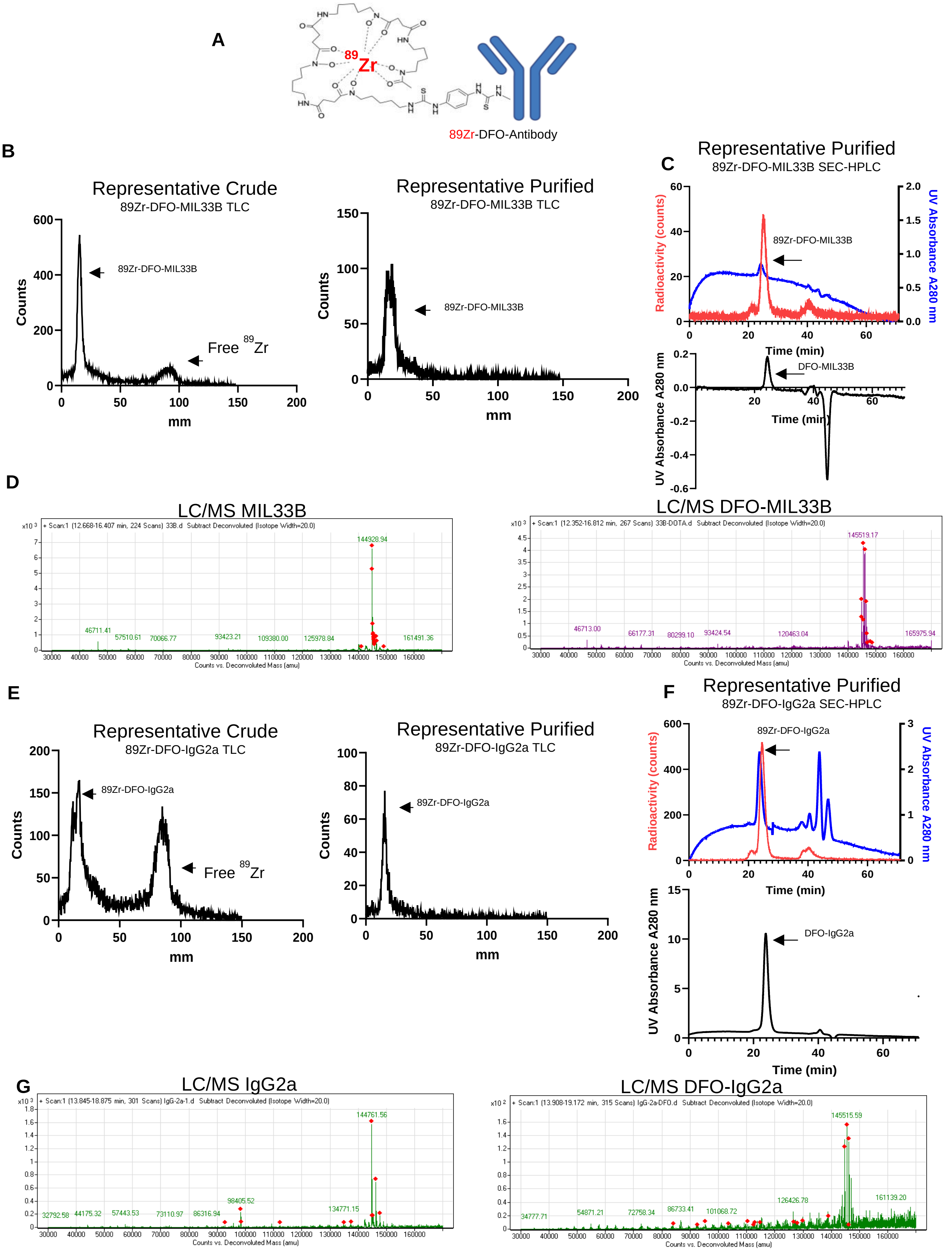
A
89Zr-DFO-Antibody
B
Representative Crude
89Zr-DFO-MIL33B TLC
89Zr-DFO-MIL33B
Free 89Zr
Counts
mm
Representative Purified
89Zr-DFO-MIL33B TLC
89Zr-DFO-MIL33B
Counts
mm
C
Representative Purified
89Zr-DFO-MIL33B SEC-HPLC
89Zr-DFO-MIL33B
Radioactivity (counts)
UV Absorbance A280 nm
Time (min)
DFO-MIL33B
UV Absorbance A280 nm
Time (min)
D
LC/MS MIL33B
144928.94
Counts vs. Deconvoluted Mass (amu)
LC/MS DFO-MIL33B
145519.17
Counts vs. Deconvoluted Mass (amu)
E
Representative Crude
89Zr-DFO-IgG2a TLC
89Zr-DFO-IgG2a
Free 89Zr
Counts
mm
Representative Purified
89Zr-DFO-IgG2a TLC
89Zr-DFO-IgG2a
Counts
mm
F
Representative Purified
89Zr-DFO-IgG2a SEC-HPLC
89Zr-DFO-IgG2a
Radioactivity (counts)
UV Absorbance A280 nm
Time (min)
DFO-IgG2a
UV Absorbance A280 nm
Time (min)
G
LC/MS IgG2a
144761.56
Counts vs. Deconvoluted Mass (amu)
LC/MS DFO-IgG2a
145515.59
Counts vs. Deconvoluted Mass (amu)

**Supplemental Figure 5: Characterization of $^{89}$Zr-DFO-MIL33B and $^{89}$Zr-DFO-IgG2a. (A) Respective antibodies were conjugated to DFO and then chelated to $^{89}$Zr**. $^{89}$Zr-DFO-MIL33B labeled with 62.7% ± 13.7% (n = 4) chelation efficiency as measured by radio-TLC (B) and when purified demonstrated > 99% purity by radio-TLC (B) and 81.7% purity when characterized by radio-SEC-HPLC (C) (n = 1). LC/MS analysis of MIL33B compared to DFO-MIL33B demonstrated a mass shift consistent with 2 chelators per antibody (D). $^{89}$Zr-IgG2a-MIL33B labeled with 54.5% ± 14.2% (n = 4) chelation efficiency as measured by radio-TLC (E) and when purified demonstrated > 99% purity by radio-TLC (E) and 86.8% purity by radio-SEC-HPLC (n = 1) (F). LC/MS analysis of IgG2a compared to DFO-IgG2a demonstrated a mass shift consistent with 2 chelators per antibody (D).

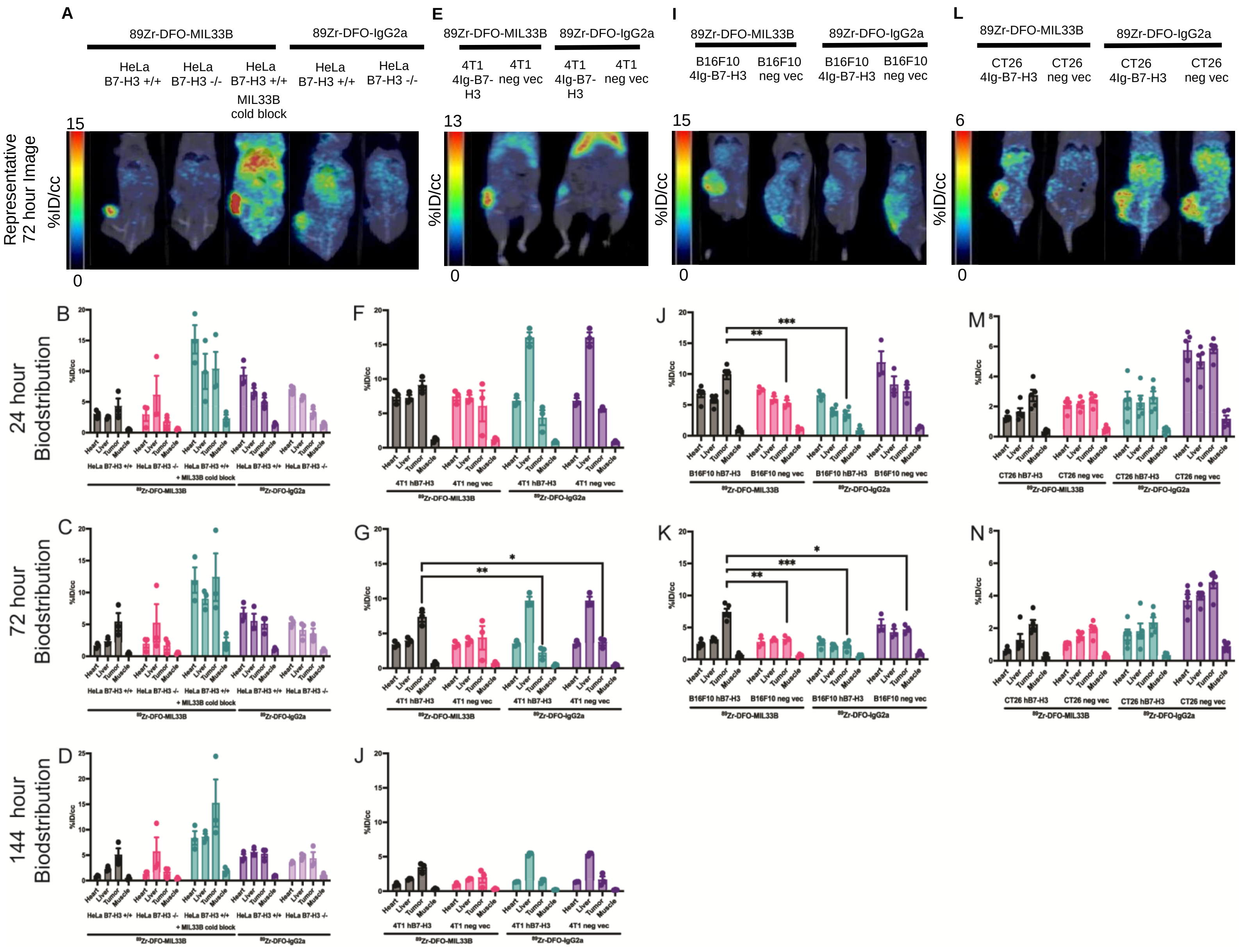

**Supplemental Figure 6: Biodistribution of $^{89}$Zr-DFO-MIL33B and $^{89}$Zr-DFO-IgG2a.** Representative 72 hour images of and 24, 72 and 144 hour biodistributions of mice imaged with $^{89}$Zr-DFO-MIL33B and $^{89}$Zr-DFO-IgG2a harboring HeLa B7-H3$^{+/+}$ and HeLa B7-H3$^{-/-}$ tumors (A-D), 4T1 4Ig-B7-H3 and 4T1 neg vec tumors (E-H), B16F10 4Ig-B7-H3 and B16F10 neg vec tumors (I-K) and CT26 4Ig-B7-H3 and neg vec tumors (L-N). (Comparisons are determined by un-paired two tailed t-tests, * = p < 0.05, ** = p < 0.01, *** p < 0.001).

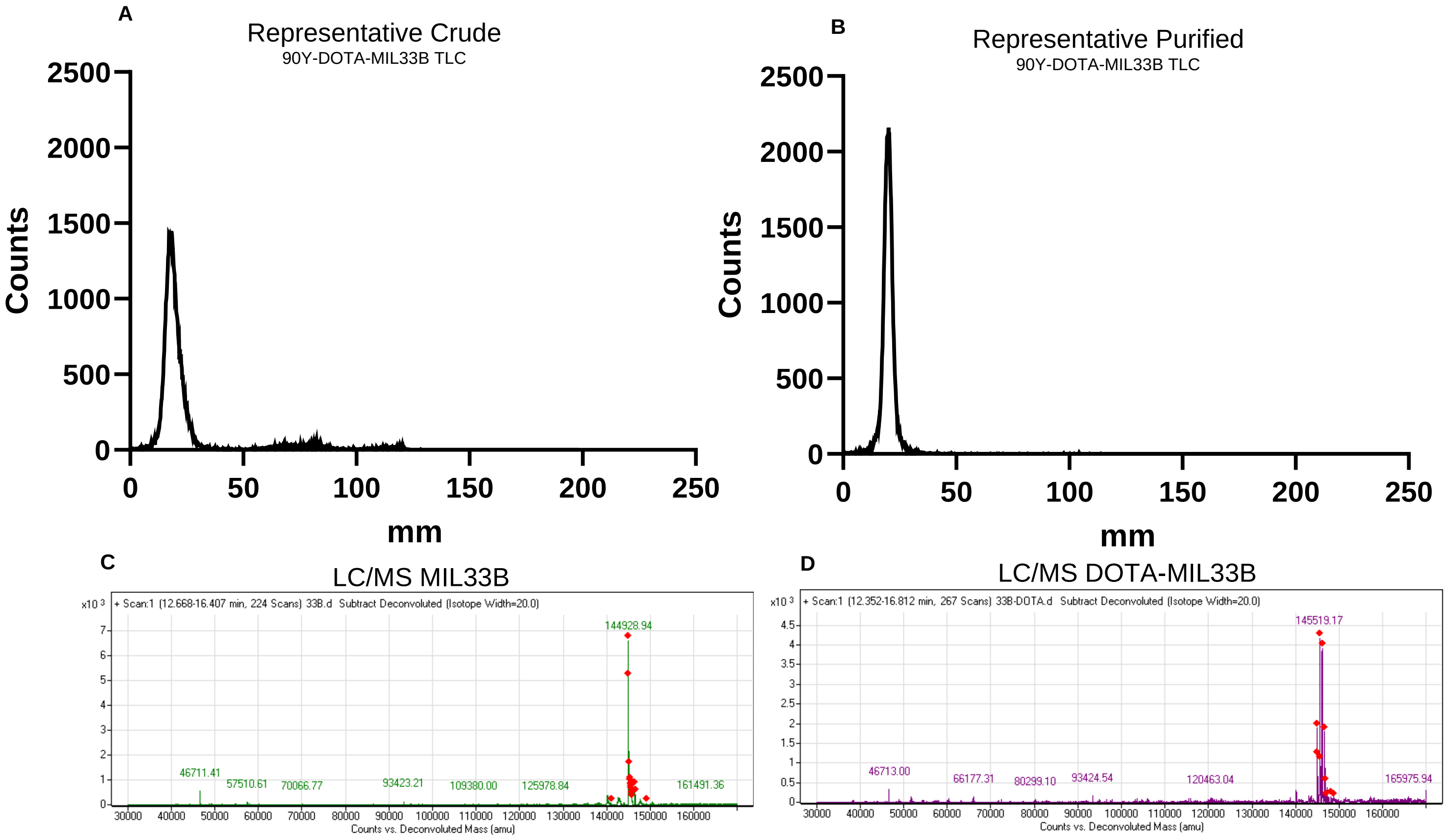


**Supplemental Figure 7: Characterization of $^{90}$Y–DOTA-MIL33B. MIL33B was conjugated to DOTA and then chelated to Yttrium-90.** DOTA-MIL33B labeled with greater than 61.9% ± 17.2% (n = 4) chelation efficiency with Yttrium-90 as determined by radio-TLC (A) and after purifying with a PD-10 column demonstrated > 99% purity by radio-TLC (B). LC/MS characterization of MIL33B (C) compared to DOTA-MIL33B (D) demonstrated a mass shift consistent with 0-1 chelators per antibody.

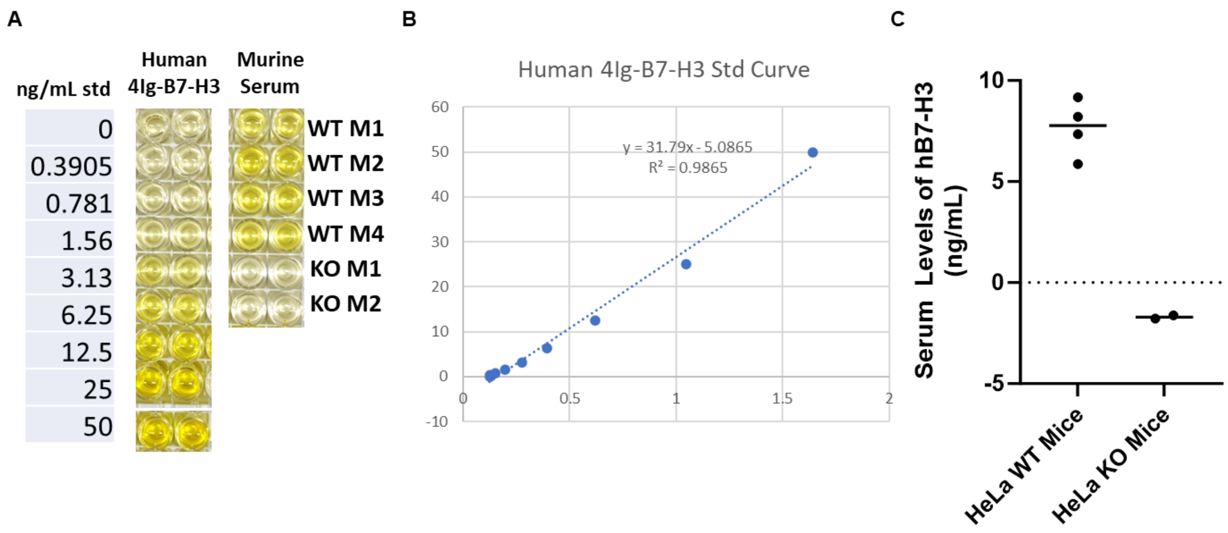


**Supplemental Figure 8: Circulating sB7-H3 is observed in HeLa xenograft serum.** Upon endpoint, HeLa WT or KO xenograft mice were subjected to a cardiac puncture and serum was isolated by centrifugation. Using a colorimetric sandwich ELISA with human B7-H3 as a standard control, serum from mice bearing WT tumors showed detectable levels of sB7-H3 (A-C).

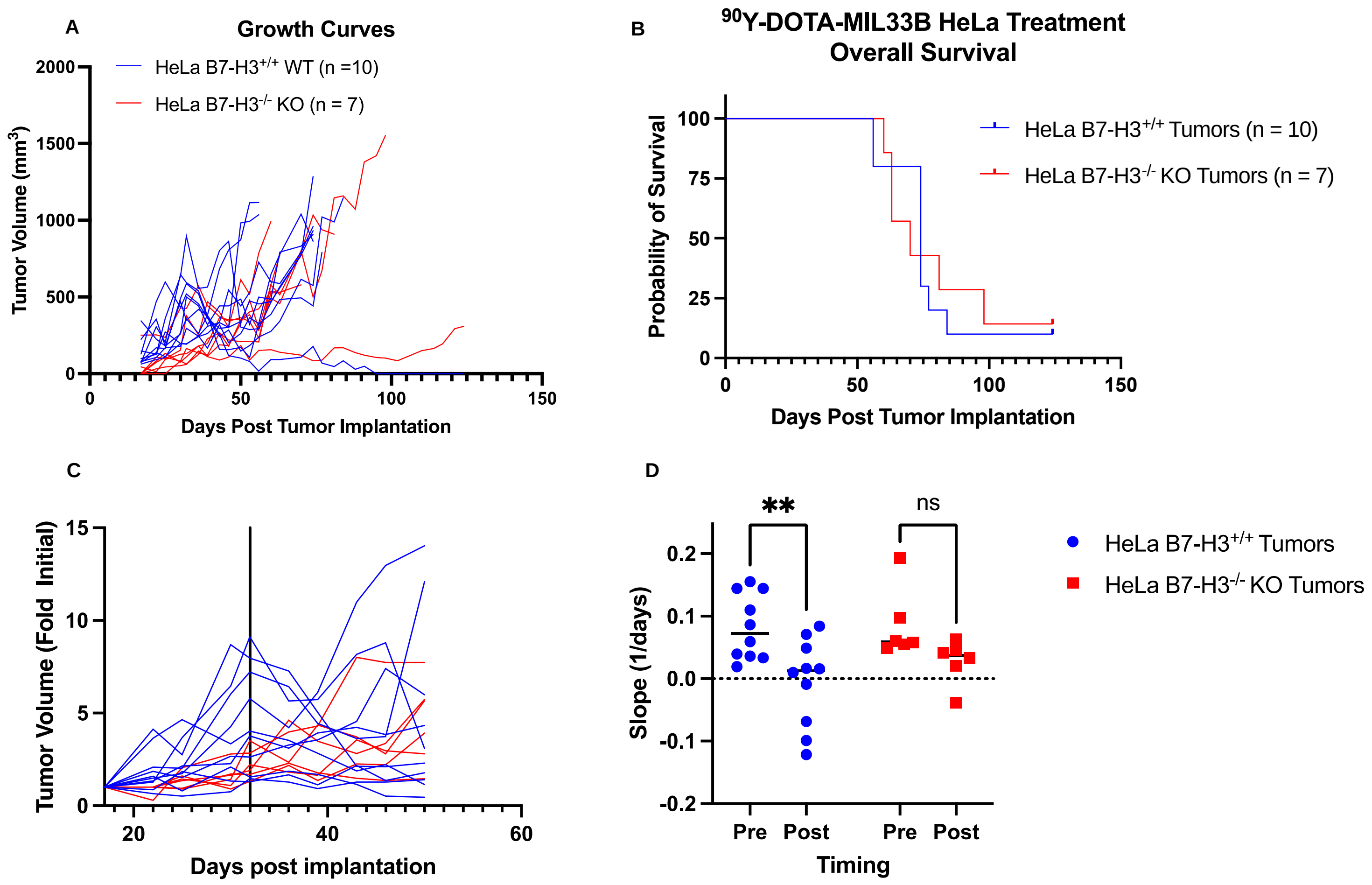


**Supplemental Figure 9:** Mice harboring HeLa B7-H3$^{+/+}$ WT tumors (n = 10) and HeLa B7-H3$^{-/-}$ KO tumors received a single i.v. dose (100 µCi) of $^{90}$Y-DOTA-MIL33B, 30 days after tumor implantation and followed for 120 days (A). There were no differences in overall survival between the groups (B), however when the slopes of the growth rates, calculated as change in tumor volume from initial palpable tumor volume, 15 days before and 15 days after treatment (treatment day is notated by the vertical line) (C), there was a significant decrease the growth rate of HeLa B7-H3$^{+/+}$ WT treated tumors (2-way ANOVA, **p = 0.0019) and no difference in growth rates of HeLa B7-H3$^{-/-}$ KO treated tumors (2-way ANOVA, p = 0.1027), demonstrating a an initial response to $^{90}$Y-DOTA-MIL33B treatment (D).

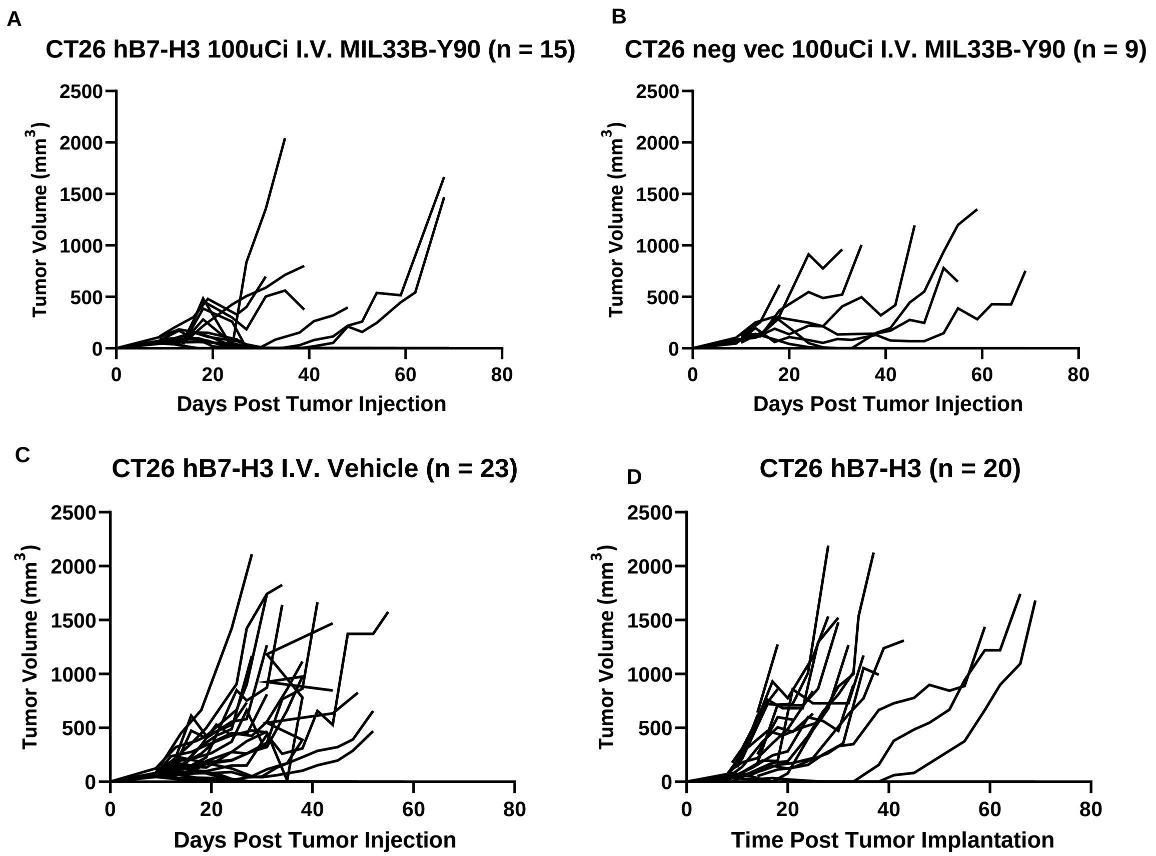


**Supplemental Figure 10: Caliper measurements of mice treated with $^{90}$Y-DOTA-MIL33B.** 15 mice harboring CT26 (h)4Ig-B7-H3 tumors received 100 µCi $^{90}$Y-DOTA-MIL33B i.v. leading to partial regression of 3 tumors and complete regression of 8 tumors (A). 9 mice harboring CT26 neg vec tumors received 100 µCi $^{90}$Y-DOTA-MIL33B i.v. leading to partial regression of 1 tumor and complete regression of one tumor (B). We also observed regression of 3 out of 23 CT26 4Ig-B7-H3 tumors that received i.v. saline (C) and spontaneous regression of 1 out of 20 untreated CT26 4Ig-B7-H3 tumors (D).

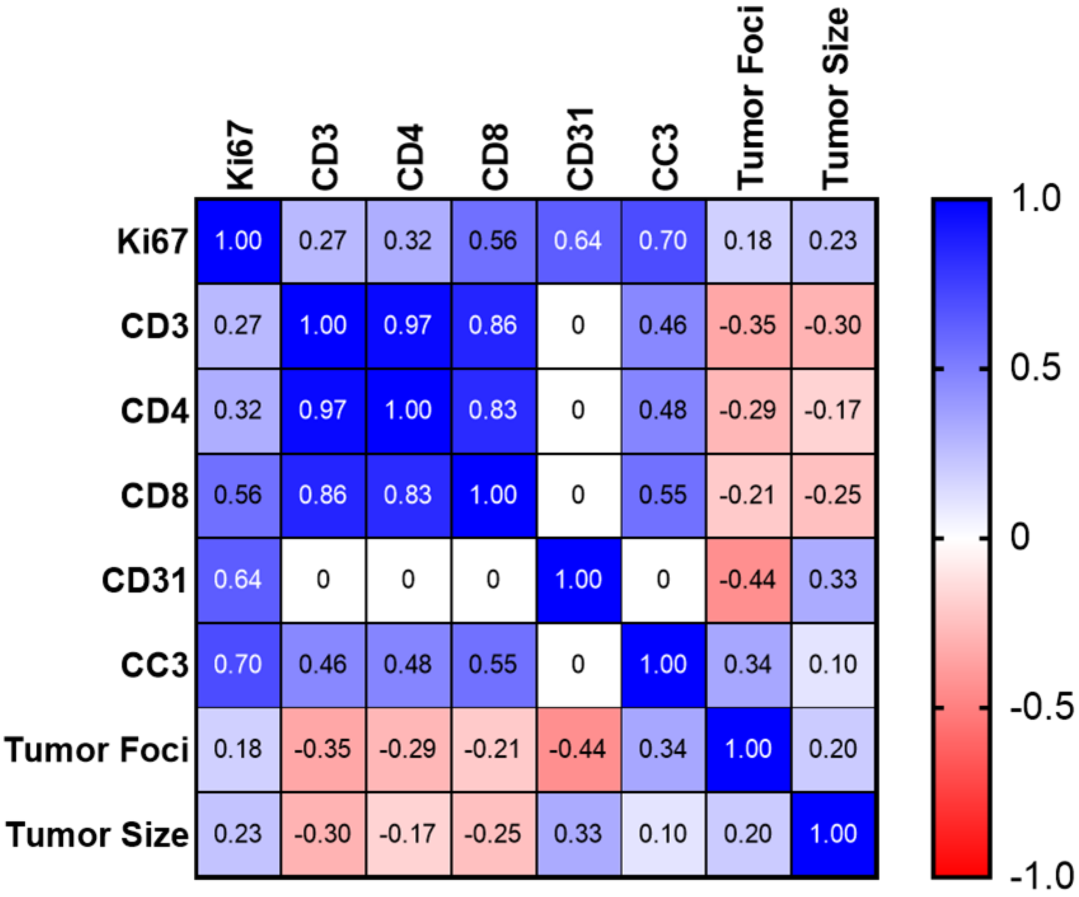


**Supplemental Figure 11: Correlation co-efficient of histology parameters indicate that adaptive immune cells correlate across samples.** Tumors were stained for the indicated markers, and then independently scored by a board-certified veterinary pathologist. Scores were then correlated across all samples to identify relevant patterns. $CD3^+$, $CD8^+$, and $CD4^+$ cells correlated with each other ($p < 0.05$, uncorrected for multiple tests).

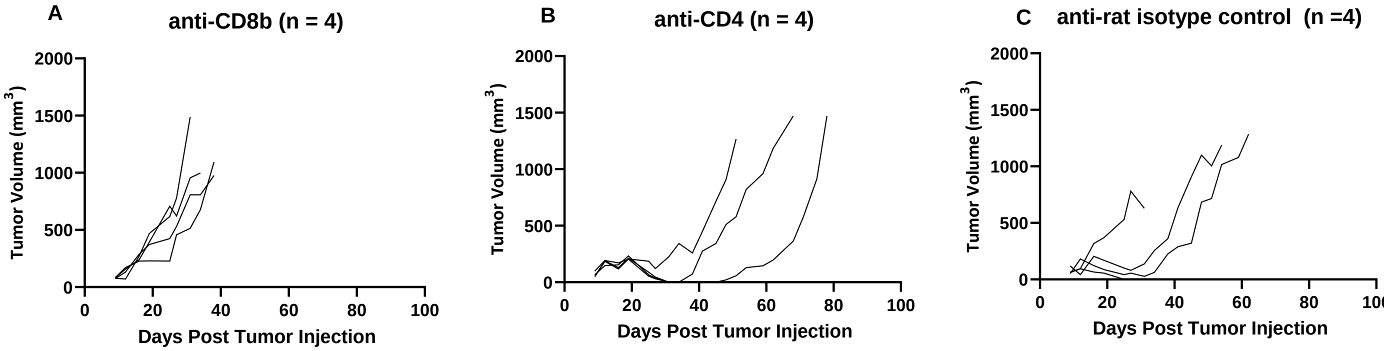


**Supplemental Figure 12: Caliper measurements of *in vivo* depletion experiment.** CT26 4Ig-B7-H3 mice treated with 100 µCi $^{90}$Y-DOTA-MIL33B i.v. after pre-treatment with either anti-CD8b-depleting, anti-CD4-depleting or anti-rat isotype control antibody. We observed no tumor regression when mice received anti-CD8b-depleting antibodies (A), tumors of 3 out of 4 mice that received anti-CD4-depleting antibodies initially regressed and 1 mouse became a long-term survivor (B). We observed similar results when treated mice received anti-rat isotype control antibody with 1 out of 4 tumors completely regressing and becoming a long-term survivor (C).

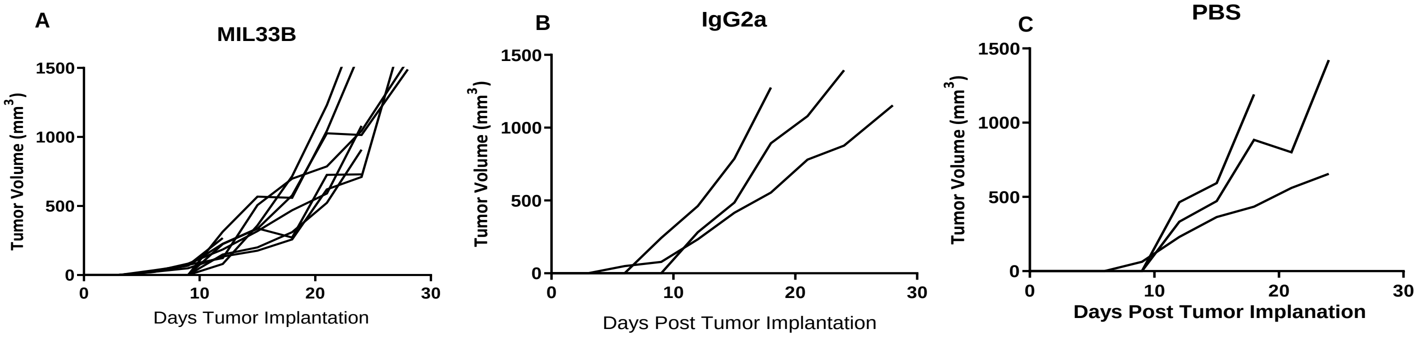


**Supplemental Figure 13: Caliper measurement of mice treated with MIL33B monotherapy compared to relevant controls.** Mice harboring CT26 4Ig-B7-H3 tumors (n = 9) received 200 µg i.p. of MIL33B every three days post tumor cell injection. We observed no tumor regression in these mice (A) or mice treated with the same dosing regimen of mouse IgG2a isotype control (n = 3) (B) or an equivalent volume of PBS (C), demonstrating no therapeutic efficacy of MIL33B as a standalone therapy in this model.
